# pOpsicle: An all-optical reporter system for synaptic vesicle recycling combining pH-sensitive fluorescent proteins with optogenetic manipulation of neuronal activity

**DOI:** 10.1101/2022.12.20.521193

**Authors:** Marius Seidenthal, Barbara Jánosi, Nils Rosenkranz, Noah Schuh, Nora Elvers, Miles Willoughby, Xinda Zhao, Alexander Gottschalk

## Abstract

pH-sensitive fluorescent proteins are widely used to study synaptic vesicle (SV) fusion and recycling. When targeted to the lumen of SVs, fluorescence of these proteins is quenched by the acidic pH. Following SV fusion, they are exposed to extracellular neutral pH, resulting in a fluorescence increase. SV fusion, recycling and acidification can thus be tracked by tagging integral SV proteins with pH-sensitive proteins. Neurotransmission is generally stimulated by electrophysiology, which is not feasible in small, intact animals, thus limiting the approach to cell culture regimes. Previous *in vivo* approaches depended on distinct (sensory) stimuli, thus limiting the addressable neuron types. To overcome these limitations, we established an all-optical approach to stimulate and visualize SV fusion and recycling. We combined distinct pH-sensitive fluorescent proteins (inserted into the SV protein synaptogyrin) and light-gated channelrhodopsins (ChRs) for optical stimulation, overcoming optical crosstalk and thus enabling an all-optical approach. We generated two different variants of the **p**H-sensitive **op**togenetic reporter of ve**sicle** recycling (pOpsicle) and tested them in cholinergic neurons of intact *Caenorhabditis elegans* nematodes. First, we combined the red fluorescent protein pHuji with the blue-light gated ChR2(H134R), and second, the green fluorescent pHluorin combined with the novel red-shifted ChR ChrimsonSA. In both cases, fluorescence increases were observed after optical stimulation. Increase and subsequent decline of fluorescence was affected by mutations of proteins involved in SV fusion and endocytosis. These results establish pOpsicle as a non-invasive, all-optical approach to investigate different steps of the SV cycle.

## 1 Introduction

Chemical synaptic transmission, the release of neurotransmitters into the synaptic cleft, depends on synaptic vesicle (SV) exocytosis (Sudhof, 2013). To efficiently sustain neurotransmitter release during phases of (high) neuronal activity, SV-associated proteins and lipids must be recycled from the plasma membrane, thus allowing to regenerate ready-to-release SVs (Gan and Watanabe, 2018; Chanaday et al., 2019). Several modes of SV recycling have been uncovered such as the classical clathrin-mediated endocytosis, activity-dependent bulk endocytosis, kiss-and-run release and the recently described ultrafast endocytosis (Heuser and Reese, 1973; Kittelmann et al., 2013; Watanabe et al., 2013a; Watanabe et al., 2013b; Morton et al., 2015; Watanabe and Boucrot, 2017; Shin et al., 2021). It is still under debate which of these processes happen under which conditions, and whether some of these represent short-cuts in the SV cycle, e.g. bypassing the endosome. Also, the exact involvement of known, and the discovery of novel, recycling factors mediating these events, is the subject of ongoing research (Gan and Watanabe, 2018; Yu et al., 2018). To study SV fusion and recycling, methods such as electron microscopy (EM), measurement of membrane capacitance and super-resolution microscopy are used. However, these methods are either limited in their temporal resolution (EM) or applicability to different neuronal cell types (membrane capacitance measurements and super-resolution microscopy) (von Gersdorff and Matthews, 1999; Yu et al., 2018; Shin et al., 2021).

Another method to indirectly visualize and quantify exocytosis and recycling of proteins is through tagging with pH-sensitive fluorescent proteins (Miesenbock et al., 1998). SVs must be acidified for refilling with neurotransmitters during the recycling process (Egashira et al., 2015; Gowrisankaran and Milosevic, 2020). For this reason, the fluorescence of pH-sensitive proteins, such as the green fluorescent pHluorin, when located on the intraluminal side of the SV membrane, is quenched by the low-pH environment (Sankaranarayanan et al., 2000). Upon SV fusion with the plasma membrane, the fluorescence increases due to exposure to the neutral extracellular medium and dequenching of the fluorophore (**Fig. 1A**). Subsequent to stimulation, the fluorescence decreases depending on the rate of endocytosis, sorting and reacidification of SVs. This way, positive or negative influences on the speed of SV retrieval can be quantified (Morton et al., 2015; Watanabe et al., 2018). Several variants of pH-sensitive fluorescent proteins have been created, covering almost the entire spectrum of visible light (Miesenbock et al., 1998; Shen et al., 2014; Liu et al., 2021). This opens the door for multiplexing with other fluorophores or optical actuators (Li and Tsien, 2012; Jackson and Burrone, 2016).

**Fig. 1.**
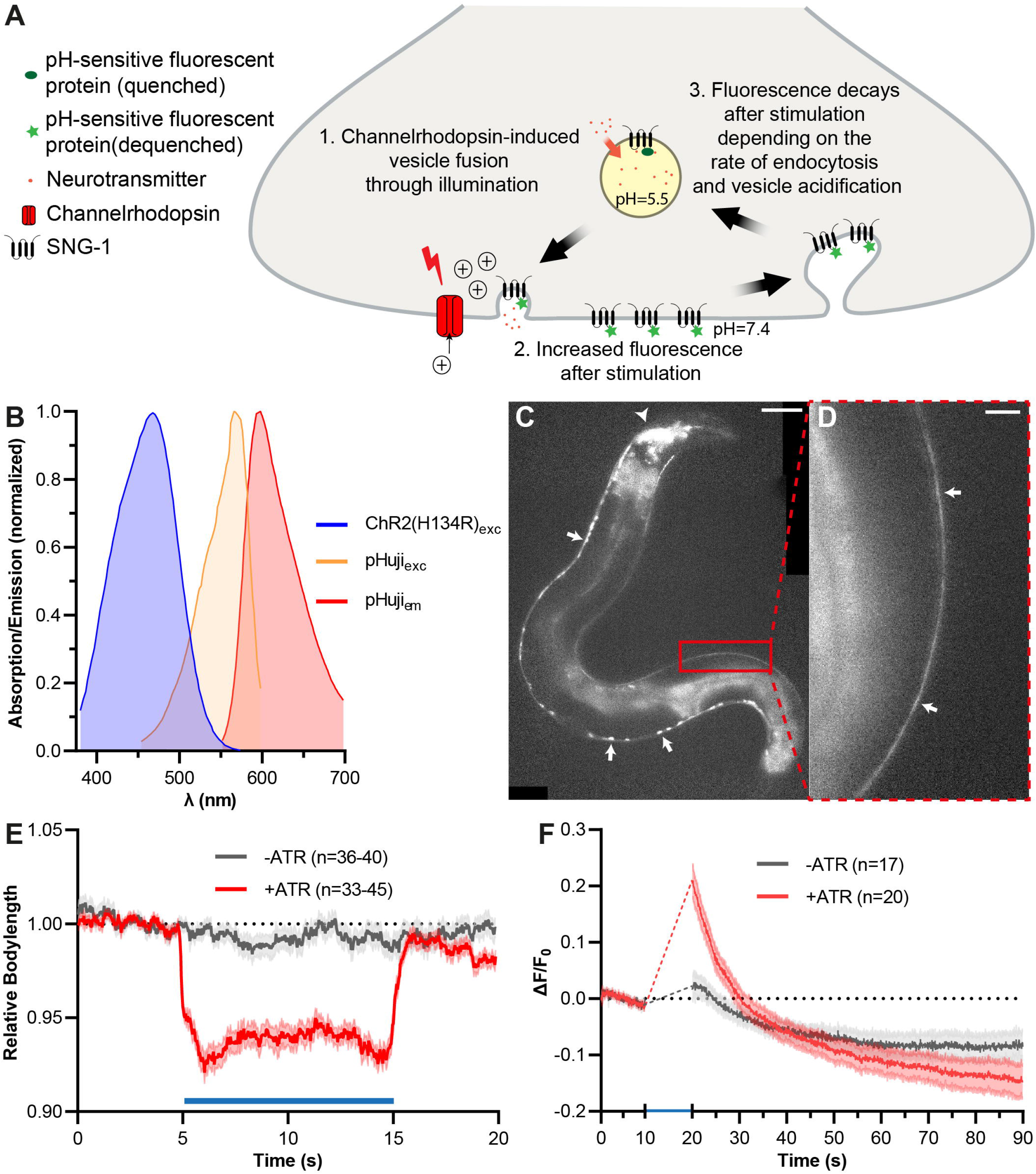
Depolarization of cholinergic motor neurons with ChR2 triggers fusion of SVs containing SNG-1::pHuji. **(A)** Schematic of the pOpsicle assay, showing the SV cycle and transfer of pH-sensitive fluorescent proteins fused to SNG-1 in response to optogenetic stimulation. **(B)** Relative excitation and emission spectra of ChR2 and pHuji, normalized to the maximum absorption/emission amplitude. **(C)** Representative image of *C. elegans* expressing SNG-1::pHuji in cholinergic neurons. Arrows: Ventral location of cell bodies of A- and B-type motor neurons. Arrowhead: cholinergic neurons within the head ganglia. 40 x magnification. Scale bar, 50 μm. **(D)** Enlarged view of the dorsal nerve cord (DNC) in (C). Arrows: fluorescent puncta, representing SV clouds and neuronal plasma membrane. Scale bar, 10 μm. **(E)** Mean relative body length (± SEM) of animals expressing ChR2 and pHuji, optionally treated with ATR (as indicated), normalized to the average body length before stimulation. A 10 s continuous light pulse (470 nm, 1 mW/mm^2^) was applied after 5 s (indicated by the blue bar). Number of animals indicated (n), accumulated from N = 3 biological replicates. **(F)** Mean (± SEM) relative change of DNC fluorescence of animals treated with and without ATR, before and after a 10 s continuous light pulse (460 nm, 0.34 mW/mm^2^), applied after 10 s (note, fluorescence of pHuji cannot be properly imaged during blue light pulse, due to photoswitching). Number of animals indicated (n), accumulated from N = 2 biological replicates.

Most studies utilizing pH-sensitive fluorescent proteins in mammalian organisms are performed using cultured neurons (Watanabe et al., 2018). *In vivo* studies are rare and usually performed in translucent non-mammalian model systems such as *Danio rerio, Drosophila melanogaster* larvae, or *Caenorhabditis elegans* (Poskanzer et al., 2003; Koudelka et al., 2016; Ventimiglia and Bargmann, 2017; Seitz and Rizzoli, 2019). Neurotransmission in these animals can be triggered by, however, labor-intensive and invasive electrophysiological stimulation. Alternatively, exposure to stimuli such as odors can be used. Yet, this is difficult to control, and limited to applications in sensory neurons (Choi et al., 2021). Thus, an all-optical solution that is not limited to certain cell types would be ideal, e.g., involving a combination of genetically encoded non-invasive tools for *in vivo* stimulation of neurons, with pH-sensitive fluorescent proteins. One possibility to manipulate neurotransmission is through transgenic expression of channelrhodopsins (ChRs), which are light-gated cation channels that can be used to depolarize neurons (Nagel et al., 2003; Boyden et al., 2005; Nagel et al., 2005; Liewald et al., 2008). Light absorption leads to isomerization of the chromophore *all-trans* retinal (ATR) and opening of the channel pore. A variety of ChRs that are activated by different wavelengths have been discovered or engineered (Guru et al., 2015; Chang, 2019). This enables multiplexing with both short- or long-wavelength absorbing fluorophores (Wabnig et al., 2015; Hawk et al., 2021; Vierock et al., 2021). In this work, we characterize two different combinations of ChRs with pH-sensitive fluorescent proteins in living *C. elegans* nematodes. We first tested pHuji, a recently described red fluorescent protein, together with the well described blue light-gated ChR2 (Nagel et al., 2005; Shen et al., 2014). This approach worked, however, not always robustly, thus we swapped both the actuator and the sensor to different excitation wavelengths. We used the recently described red-light activated ChR ChrimsonSA and the well-established green fluorescent pHluorin (Miesenbock et al., 1998; Oda et al., 2018; Seidenthal et al., 2022). Using this combination, we could stimulate and visualize SV exo- and endocytosis in an all-optical, non-invasive manner *in vivo*. We termed this approach the **p**H-sensitive **op**togenetic reporter of ve**sicle** recycling (pOpsicle). We tested the pOpsicle method in cholinergic motor neurons and in the glutamatergic/tyraminergic interneuron RIM. pOpsicle should be applicable to most neuronal cell types and is, to our knowledge, the only all-optical approach to study SV recycling using ChRs and pH-sensitive fluorescent proteins in living animals to date. Our approach expands the possibilities to study SV recycling at the *C. elegans* neuromuscular junction (NMJ) which previously could only be done by indirect measurement of postsynaptic effects using electrophysiology, Ca^2+^ imaging, or by (non-)time-resolved electron microscopy (Liewald et al., 2008; Kittelmann et al., 2013; Wabnig et al., 2015; Steuer Costa et al., 2017; Yu et al., 2018).

## 2 Materials and methods

### 2.1 Molecular biology

For the expression of SNG-1 fusion constructs and channelrhodopsins in *C. elegans*, the *punc-17* promotor (cholinergic motor neurons) and a short version of the *ptdc-1* promotor (RIM interneurons) were used. **pcDNA3-SypHluorin 4x (S4x)** was a gift from Stephen Heinemann & Yongling Zhu (Addgene plasmid #37005; http://n2t.net/addgene:37005; RRID: Addgene_37005). **pJB14** (*TOPO vector::2xpHluorin*) was generated using the TOPO cloning kit (Thermo Fisher Scientific Inc., USA) by amplifying two copies of pHluorin cDNA from the pcDNA3-SypHluorin 4x (S4x) plasmid with primers oBJ51 (5’-ATATCGAACCGTCTTCAGATATGGATCTAGCCACC-3’) and oBJ62 (5’-TATATTCGCCGTCTTCTCCACCGCATGTGATTCGAGCTCC-3’). **pJB10** (*punc-17::sng-1::unc-54-3’UTR*) was generated through Gibson assembly by digesting pRM348 with *BmtI* and *BsiWI* (*punc-17* and backbone), by amplifying pAG52 (*sng-1*) with primers oBJ58 (5’-TCAGGAGGACCCTTGGCTAGATGGAGAACGTGCGTGCTTATG-3’) and oBJ59 (5’-ATGACTCGAGCTAATAACCATATCCTTCCGACTGAG-3’) and by amplifying pAH03 (*unc-54-3’UTR*) with oBJ60 (5’-ATATGGTTATTAGCTCGAGTCATGGTCGACAAG-3’) and oBJ61 (5’-AAACGCGCGAGACGAAAGGGCCCAAACAGTTATGTTTGGTATATTGGG-3’). **pJB11** (*punc-17:: sng-1::2xpHluorin::unc-54-3’UTR*) was generated by digestion of pJB10 and pJB14 with *BbsI* and subsequent ligation to introduce two copies of pHluorin cDNA into the sequence encoding the first intraluminal loop of SNG-1. **pDisplay-pHuji** was a gift from Robert Campbell (Addgene plasmid #61556; http://n2t.net/addgene:61556; RRID: Addgene_61556). pDisplay-pHuji was amplified using primers oBJ104 (5’-GCAGAAGAAAACCATGGGCTG-3’) and oBJ105 (5’-CAGCCCATGGTTTTCTTCTGC-3’) to remove the *BbsI* restriction site to generate **pJB24**. **pJB25** (*punc-17::sng-1::pHuji::unc-54-3’UTR*) was generated via Gibson assembly by digesting pJB10 with *BbsI* and amplifying pJB24 with primers oBJ107 (5’-ATATCGAAAAGTCTTCAGGTGGAGGTGGAAGTATGGTGAGCAAGGGCGAG-3’) and oBJ108 (5’-TATATTCGCCGTCTTCGGTGGAGGTGGAAGTCTTGTACAGCTCGTCCATG-3’) which contain the sequence for a GGGGS linker to add in front of the coding region of pHuji. **pJB26** (*punc-17::ChR2(H134R)::myc*) was generated by amplifying *ChR2(H134R)::myc* using primers oBJ113 (5’-GAACGCTAGCACCACTAGATCCATCTAGAG-3’) and oBJ114 (5’-GCATGCTAGCCACCAGACAAGTTGGTAA-3’) which was introduced into pRM348 by restriction digest with *NheI* and subsequent ligation. **pMSE01** (*punc-17::ChrimsonSA::unc-54-3’UTR*) was generated by amplifying pDV07 (*punc-17::ChrimsonWT::unc-54-3’UTR*) with oMSE16 (5’-CGAGTGGCTGCTGGCTTGCCCCGTGAT-3’) and oMSE017 (5’-ATCACGGGGCAAGCCAGCAGCCACTCG-3’) to introduce the point mutation (S169A). **pMSE23** (*ptdc-1s::ChrimsonSA::unc-54-3’UTR*) was generated by amplifying *ptdc-1s* from pXY07 (*ptdc-1s::GFP*) with primers oMSE105 (5’-TCCCGGCCGCCATGGCCGCGATTTCTGTATGAGCCGCCCG-3’) and oMSE106 (5’-AAAGACTTTCGATGAATTACTTGGGCGGTCCTGAAAAATG-3’), amplifying the ChrimsonSA backbone from pMSE01 with oMSE107 (5’-CATTTTTCAGGACCGCCCAAGTAATTCATCGAAAGTCTTTCTATTTTCCGCATCTCTTGTTCAAGGGATTGG-3’) and oMSE108 (5’-CGGGCGGCTCATACAGAAATCGCGGCCATGGCGGCCG-3’) and combined the fragments using Gibson assembly. **pMSE24** (*ptdc-1s::sng-1::pHluorin::unc-54-3’UTR*) was generated by amplifying the *sng-1::pHluorin* backbone from pBJ11 with oMSE114 (5’-AGGGTCGACCATGACTCGAGCTAATAACCATATCCTTC-3’) and oMSE115 (5’-GTAATTCATCGAAAGTCTTTCTATTTTCCGCATCTCTTGTTCAAGGGATTGG-3’) and with oMSE108 and oMSE113 (5’-GAAGGATATGGTTATTAGCTCGAGTCATGGTCGACCCT-3’) and fusing these fragments with the *ptdc-1s* using Gibson assembly.

### 2.2 Cultivation of *C. elegans*

*C. elegans* strains were kept under standard conditions on nematode growth medium (NGM) plates seeded with the *Escherichia coli* strain OP50, obtained from the *Caenorhabditis* Genetics Center (CGC, University of Minnesota, USA), at 20°C (Brenner, 1974). The N2 Bristol strain was provided by the CGC and used as wild type. Transgenic animals were generated as described previously (Fire, 1986). An overview of transgenic and mutant strains used or generated in this work can be found in Table 1. For experiments, well-fed L4 larvae were picked ~18 h before the assays. For RIM experiments, only animals showing marker fluorescence were used. Animals were supplemented with ATR (Sigma-Aldrich, USA) by adding 100 μl OP50 containing 200 μM ATR to 10 ml NGM agar dishes. Experiments were performed on at least two separate days with animals picked from different plates.

**Table 1:** *C. elegans* strains.

| Strain | Genotype | Source |
| --- | --- | --- |
| ZX214 | *snt-1(md290)* | CGC (Jorgensen et al., 1995) |
| ZX307 | *snb-1(md247)* | CGC (Miller et al., 1996) |
| ZX451 | *unc-57(e406)* | CGC (Brenner, 1974) |
| ZX1629 | *unc-26(s1710)* | CGC (Harris et al., 2000) |
| ZX2835 | *sng-1(ok234); zIs138[punc-17::sng-1-pHuji, punc-17::Chr2(H134R)::myc, pmyo-2::CFP]* | This study |
| ZX2836 | *unc-26(s1710); sng-1(ok234); zIs138[punc-17::sng-1-pHuji, punc-17::Chr2(H134R)::myc, pmyo-2::CFP]* | This study |
| ZX2837 | *snt-1(md290); sng-1(ok234); zIs138[punc-17::sng-1-pHuji, punc-17::Chr2(H134R)::myc, pmyo-2::CFP]* | This study |
| ZX2838 | *unc-57(e406); sng-1(ok234); zIs138[punc-17::sng-1-pHuji, punc-17::Chr2(H134R)::myc, pmyo-2::CFP]* | This study |
| ZX2850 | *snb-1(md247); sng-1(ok234); zIs138[punc-17::sng-1-pHuji, punc-17::Chr2(H134R)::myc, pmyo-2::CFP]* | This study |
| ZX3197 | *zIs152[punc-17::Chrimson(S169A); punc-17::pHluorin; pmyo-2::mCherry]* | This study |
| ZX3217 | *unc-26(s1710); zIs152[punc-17::Chrimson(S169A); punc-17::pHluorin; pmyo-2::mCherry]* | This study |
| ZX3218 | *snb-1(md247); zxIs152[punc-17::Chrimson(S169A); punc-17::pHluorin; pmyo-2::mCherry]* | This study |
| ZX3254 | *unc-57(e406); zxIs152[punc-17::Chrimson(S169A); punc-17::pHluorin; pmyo-2::mCherry]* | This study |
| ZX3402 | *snt-1(md290); zxIs152[punc-17::Chrimson(S169A); punc-17::pHluorin; pmyo-2::mCherry]* | This study |
| ZX3422 | *zxEx1418[pmyo-2::mCherry; ptdc-1s::ChrimsonSA; ptdc-1s::SNG-1::pHluorin]* | This study |

### 2.3 Measurement of *C. elegans* body length

Body length assays were performed as described previously (Liewald et al., 2008; Seidenthal et al., 2022). Briefly, ChR2(H134R) was activated using a 450-490 nm bandpass excitation filter at 1 mW/mm^2^ light intensity. ChrimsonSA was stimulated using light from a 50 W HBO lamp filtered through a 590-650 nm filter, and adjusted to 1 mW/mm^2^ light intensity. Brightfield light was filtered with a 665-715 nm filter to avoid unwanted activation of channelrhodopsins. Videos of single animals were acquired and then analyzed using the WormRuler software (Seidenthal et al., 2022). Body length of each worm was normalized to the 5 s period before stimulation and values >120 % or < 80% of the initial body length were discarded as these are biomechanically impossible and result from artifacts in the background correction.

### 2.4 Measurement of crawling speed and reversals using the multi-work-tracker (MWT)

Videos of crawling animals were acquired as described previously (Vettkötter et al., 2022) and crawling speed measured using the MWT setup (Swierczek et al., 2011). Animals were washed three times with M9 buffer to remove OP50 bacteria. They were then transferred to unseeded NGM plates and kept in darkness for 15 minutes. A light stimulus was applied using a custom-build LED ring (Alustar 3W 30°, ledxon, 623 nm) which was controlled by an Arduino Uno (Arduino, Italy) device running a custom-written Arduino script. Videos were acquired using a high-resolution camera (Falcon 4M30, DALSA) and crawling speed of single animals as well as reversal count (in bins of 10 s) were extracted using ‘Choreography’ software (Swierczek et al., 2011) and summarized using a custom Python script.

### 2.5 Microscopy and imaging

For fluorescence imaging, animals were placed upon 7 % agarose pads in M9 buffer. Animals were immobilized using a 20 mM Levamisole-hydrochloride (Sigma-Aldrich, USA) solution in M9 and visualized on an Axio Observer Z1 microscope (Zeiss, Germany) equipped with a 100 x oil objective. Fluorescent proteins and channelrhodopsins were excited using a 460 nm and a 590 nm LED system (Lumen 100, Prior Scientific, UK) coupled via a beamsplitter. pHuji and ChR2(H134R) were excited using a double band pass filter (460 – 500 nm, 570 – 600 nm) combined with a 605 nm beam splitter (AHF Analysentechnik, Germany). 460 nm LED light to stimulate ChR2(H134R) was set to 340 μW/mm^2^ intensity. pHuji fluorescence was filtered using a 615 – 680 nm emission filter and visualized using an EMCCD camera (Evolve 512 Delta, Teledyne Photometrics, USA). pHluorin and ChrimsonSA were excited using a 450 - 490 nm/ 555 – 590 nm double band pass filter combined with a GFP/mCherry beamsplitter (AHF Analysentechnik, Germany). 590 nm LED light intensity to stimulate ChrimsonSA was set to 40 μW/mm^2^. pHluorin fluorescence was filtered using a 502.5 – 537.5 nm band pass emission filter and visualized using a sCMOS camera (Kinetix 22, Teledyne Photometrics, USA). The dorsal nerve cord (DNC) was visualized using the basal pHuji or pHluorin fluorescence. For cholinergic neurons, the a region in the posterior third of the animal was imaged, where an abundance of synaptic puncta can be found. For RIM experiments, fluorescent neuronal extensions in the head region were visualized. Videos were captured using the μManager v.1.4.22 software (Edelstein et al., 2014). pHuji - pOpsicle experiments were performed with 50 ms exposure time, pHluorin - pOpsicle experiments with 200 ms exposure. Stimulation of channelrhodopsins was triggered using a custom written *Autohotkey* script to activate and deactivate LEDs. Representative images displaying entire worms were acquired using a 40 x oil objective and stitched together using the ImageJ *Stitching* Plugin (Preibisch et al., 2009). The representative image of RIM neurons (**Fig. 6A**) was made using the *Z Project* function to generate a projection of slices acquired throughout the head region.

### 2.6 Quantification of fluorescence

Example images were processed, and fluorescence was quantified using ImageJ v1.53 (Schindelin et al., 2012). A region of interest (ROI) was placed on the DNC or RIM neuron using the *Segmented Line* tool. Pixel width of the line was adjusted according to the width of the fluorescent signal. A background ROI was set in close proximity to the imaging ROI, inside the worm (but avoiding gut autofluorescence) and fluorescence was quantified using the *Multi Measure* function. XY-drift was corrected using the *Template Matching* ImageJ plugin, if necessary. Animals that moved excessively or drifted in the focal plane were discarded. Fluorescence was normalized to the average fluorescence before stimulation (F_0_) to compare different animals:

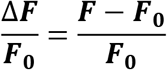

A custom written python script was used for background subtraction, normalization and (if needed) filtering of animals according to whether they show a strong response during stimulation (available on GitHub^1^). For this, the maximum background corrected fluorescence during stimulation was calculated (as a moving average of 1 s). If this was higher than the average background corrected fluorescence before the stimulation + 3 * standard deviation (of background corrected fluorescence before stimulation), the animal was counted as a strong responder (adapted from Choi et al., 2021); see **Fig. S3A** for example traces that fit or do not fit these parameters; animals not fitting the cut-off showed no discernable light-evoked effect on DNC fluorescence. Also, animals that showed an increase of the fluorescence after the end of the stimulus, or animals showing spontaneous events, were excluded. These measures were necessary for the calculation of fluorescence rise and decay kinetics, since data from ‘non-responders’ could not be properly fitted. Fluorescence was not corrected for bleaching since the measured background fluorescence bleached with a similar rate as the fluorescence in the DNC. Thus, subtraction of background fluorescence was sufficient to correct for bleaching. Attempts to further correct for bleaching led to a progressive deviation towards the end of the acquisition. Fluorescence signal increases in the pOpsicle assay were calculated using the mean of the normalized fluorescence at the first second after stimulation (± SEM) for pHuji experiments or the mean of seconds 15 to 20 (± SEM) for pHluorin experiments. Regression analysis, to calculate the rate of fluorescence rise and decay, was performed using Graphpad Prism 9.4.1. One-phase exponential association (1) and decay (2) equations were fitted to the timepoints during and after stimulation and the time constants τ calculated for each animal:

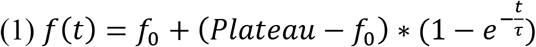

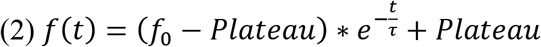

t: time (in seconds)
*f*_0_: value of *f*(*t*) at t = 0
Plateau: value of *f*(*t*) at t = ∞
τ: time constant (in seconds; higher τ values indicate a slower rise or decay)

As above, each fit was inspected. Individual fits that showed no increase during stimulation were discarded from analysis as they could be fitted properly (**Fig. S3A**). Similarly, data sets that displayed an increase rather than a decay after stimulation were also discarded.

### 2.7 *C. elegans* Primary neuronal cell culture

For the preparation of *C. elegans* primary cell culture, established protocols were adapted and modified (Christensen et al., 2002; Strange et al., 2007). Gravid adult worms were grown on enriched peptone plates with nystatin (NEP agar) seeded with Na22 *E. coli* (CGC; Zhang et al., 2011). Worms were washed off the plate using double-distilled water (ddH_2_O) and transferred to 15 ml centrifuge tubes. 2 ml of household bleach as well as 1 ml of 5 M NaOH solution were added to 7 ml of worm suspension. The solution was vortexed for at least five minutes at maximum speed to get rid of adult worm bodies. All the following steps were performed under a sterile workbench. The solution, now containing only eggs, was centrifuged at 500 g for one minute. Excess liquid was removed, and the pellet was resuspended in ddH_2O_. Washing was repeated three times. The egg pellet was resuspended in 500 μl freshly thawed chitinase (1 U/ml, Sigma-Aldrich, USA) and transferred to a 1.5 ml tube. The tube was placed into a shaker for 90 minutes at room temperature to digest the chitin shell of the eggs. The chitinase reaction was stopped with 800 μl L-15 full medium (Gibco, USA) containing 10 % fetal calf serum (FCS) as well as Pen/Strep (50 U/ml penicillin + 50 μg/ml streptomycin; Sigma-Aldrich, USA). After centrifugation at 900 g, excess liquid was discarded, and the pellet was resuspended in 500 μl L-15 full medium. Using a 2 ml syringe with an 18-gauge needle, the solution was aspirated and released back into the tube 15 to 20 times to dissociate the cells. After dissociation, 1 ml L-15 full medium was added to the tube and taken up into the syringe. With the cell solution inside the syringe, the needle was replaced by a 5 μm filter (Millipore, Germany). The solution was released through the filter into a fresh tube. The original tube was refilled with 1 ml L-15 full medium, the filter was replaced by the needle and the procedure was repeated to release the solution into another tube. This was repeated four to six times, depending on the initial number of eggs (more eggs = less repetitions). Filtered cell solutions were centrifuged at 900 g and most of the supernatant was discarded. Cell pellets were resuspended in the remaining medium and pooled. 500 to 1000 μl L-15 full medium were added to the suspension and the solution was seeded on 1 - 2 peanut lectin (Sigma-Aldrich, USA) coated glass bottom petri dishes (ibidi, Germany). Petri dishes were filled with 1 ml L-15 full medium and cells were allowed to adhere for 24 hours in a 20 °C sterile incubator (Memmert, Germany) before exchanging the medium. Treatment with ATR was performed after two days. Medium was replaced with a 10 μM ATR solution in L15 full medium and the cells incubated for at least 12 h at 20 °C. Prior to measurement, cells were washed two times with L-15 full medium without ATR. Neurons were imaged three to four days after seeding. The filter setup was identical to the one used to visualize pHluorin in whole animals, yet neurons were visualized using a 40 x objective. Buffers used to either de-quench or quench pHluorin fluorescence were adapted from Dittman and Kaplan (2006) and added manually by pipetting 1 ml of the respective solution onto the petri dish and then removing the same amount of liquid. Before starting a new acquisition, cells were washed three times using the control saline buffer (pH 7.4). Fluorescence was quantified as in living animals. A ROI was set on top of neurite extensions using the ImageJ “Segmented line” tool with a background ROI set in proximity.

### 2.8 Statistical Analysis

Statistical analysis and plotting of graphs were done using Graphpad Prism 9.4.1. Unpaired *t*-Tests were performed if two normally distributed datasets were compared or one-way ANOVAs using the Dunnett’s correction for multiple comparisons for three or more datasets. Fluorescence rise and decay constants were compared using the Mann-Whitney test (for two datasets) or the Kruskal-Wallis tests with Dunn’s correction for multiple comparisons (three or more datasets).

## 3 Results

### 3.1 Stimulation of neurotransmission using ChR2 triggers fusion of SVs containing synaptogyrin-pHuji

To enable an all-optical method for analysis of SV exo- and endocytosis, we used proteins with presumably low optical crosstalk. Specifically, we tested the recently described red fluorescent pHuji protein which shows a 22-fold increase of fluorescence when transferred from intravesicular to extracellular pH (Shen et al., 2014). pHuji was combined with the blue light-gated ChR2(H134R; hereafter called ChR2) for stimulation of neurotransmission (Nagel et al., 2005). The spectral overlap of the excitation spectra of pHuji and ChR2 is minimal (Azimi Hashemi et al., 2014; Shen et al., 2014; Lambert, 2019; **Fig. 1B**). To use pH-sensitive proteins as reporters for exo- and endocytosis, they must be targeted to the acidic intraluminal side of the SV membrane (Miesenbock et al., 1998; Ventimiglia and Bargmann, 2017; Li et al., 2022). Typically, the utilized reporters are either fused to the vesicular glutamate transporter or inserted into loops of the tetraspan membrane protein synaptophysin (Voglmaier et al., 2006; Luo et al., 2021). Since in *C. elegans* proteins can be targeted to e.g. cholinergic or GABAergic motor neurons using specific promoters, we chose the ubiquitous (in neurons) integral SV membrane protein synaptogyrin-1 (SNG-1), which is closely related to synaptophysin (however, *C. elegans* SPH-1 is not expressed in neurons), to target pH-sensitive fluorescent proteins (Zhao and Nonet, 2001; Abraham et al., 2006). SNG-1 has previously been used to target proteins to SVs (Liu et al., 2019; Vettkötter et al., 2022). We inserted pHuji into the second intraluminal loop of SNG-1, which should expose it to the acidic inside of SVs (**Fig. 1A**). This construct was co-expressed with ChR2 in cholinergic motor neurons using the promotor of *unc-17* which encodes the vesicular acetylcholine transporter (Alfonso et al., 1993; Miller et al., 1996; Liewald et al., 2008). SNG-1::pHuji could be observed throughout the entire cholinergic nervous system, including the nerve ring, as well as the ventral and dorsal nerve cords (VNC/DNC; **Fig. 1C**). It also localizes to neuronal extensions of cholinergic DA and DB neurons innervating dorsal muscle cells (**Fig. 1D**). Fluorescent puncta in the DNC indicate accumulation of SNG-1::pHuji at NMJs (Sieburth et al., 2005). Next, we analyzed whether neurotransmission can be activated in these animals using ChR2. Stimulation of cholinergic motor neurons leads to a reduction in body length through acetylcholine release which activates muscle cell contraction (Liewald et al., 2008). This was the case, as blue light caused a decrease of body length in animals that were grown in the presence of *all-trans retinal* (ATR; **Fig. 1E**), while controls without ATR did not alter their body length upon illumination. We therefore investigated whether stimulation of cholinergic motor neurons leads to exocytosis and thus to an increase in the fluorescence of SNG-1::pHuji. pHuji fluorescence was quantified in the DNC towards the posterior part of the animal due to the abundance of NMJs in this body region (Sieburth et al., 2005; **Fig. 1D, F**). Since pHuji fluorescence was relatively dim, we were unable to quantify fluorescence during the stimulation period: Blue light increased the autofluorescence within the animal, thus erroneously increasing the measured pHuji signal. Alternating light protocols also failed, likely due to the photoswitching nature of pHuji (Liu et al., 2021). However, it was possible to compare the fluorescence before and after stimulation. Animals treated with ATR showed a significantly increased fluorescence after stimulation (19 ± 3 % ΔF/F_0_), compared to animals without ATR (2 ± 2 %, ***p < 0.001; **Fig. 1F**). This increase gradually declined towards the fluorescence level in animals grown without ATR. The decline is most likely an indicator for recycling of externalized SNG-1::pHuji (and thus of SVs). We termed the pHuji-ChR2 combination ‘red pOpsicle’.

### 3.2 pHuji fluorescent signal increase is an indicator for SV exocytosis

To determine whether the pOpsicle assay is capable of efficiently reporting SV exo- and endocytosis, we crossed the ChR2 and SNG-1::pHuji expressing transgene into mutants known to affect proteins involved in the SV cycle (**Fig. 2**). Synaptobrevin-1 (SNB-1) is an essential vesicular soluble N-ethylmaleimide-sensitive-factor attachment receptor (v-SNARE) and thus involved in SV fusion with the plasma membrane at active zones (Nonet et al., 1998; Liu et al., 2018). Reduction-of-function mutants show reduced acetylcholine release and post-synaptic currents (Nonet et al., 1998). In comparison to wild type animals, *snb-1(md247)* mutant animals showed a significantly lower fluorescence increase after stimulation with blue light (wild type: 15 ± 2 %, *snb-1(md247)*: 2 ± 2 %; **Fig. 2A, B**), as expected, because lower amounts of SVs containing SNG-1::pHuji are exocytosed due to the defective SNARE complex. Next, we examined whether the rate of the fluorescence decay after stimulation depended on the rate of SV recycling. We thus crossed the pHuji expressing transgene into mutants lacking the established SV recycling factors synaptojanin-1/UNC-26, endophilin-1/UNC-57 and synaptotagmin-1/SNT-1. These proteins are involved in numerous processes at different steps in SV recycling, such as membrane bending and clathrin-uncoating (Jorgensen et al., 1995; Harris et al., 2000; Schuske et al., 2003; Kittelmann et al., 2013; Watanabe et al., 2018; Yu et al., 2018; Mochida, 2022). Notably, SNT-1 is also the primary sensor of calcium for exocytosis but independent from this also regulates endocytosis, as an adapter complex 2 (AP2) binding site in the plasma membrane (Poskanzer et al., 2003; Yu et al., 2013; Mochida, 2022). Both, *unc-26(s1710)* and *unc-57(e406)* knockout mutants, displayed a higher increase in pHuji fluorescence than wild type animals (**Fig. 2C:** wild type 16 ± 3 %, *unc-26(s1710)* 30 ± 5 %, *p < 0.05; **Fig. 2D:** wild type 16 ± 3 %, *unc-57(e406)* 30 ± 4 %, *p < 0.05). This was surprising since both mutants were previously shown to be depleted of ready-to-release SVs and should therefore not be able to externalize as many SVs as wild type animals (Harris et al., 2000; Schuske et al., 2003). Possibly, this could be due to reduced recycling during the stimulation period, thus leading to an accumulation of SNG-1 on the plasma membrane. Nevertheless, both mutants showed prolonged increased fluorescence during acquisition compared to wild type. *snt-1(md290)* knockout mutants displayed a similar signal increase and decay trend as wild type (**Fig. 2E:** wild type 14 ± 3 %, *snt-1(md290)* 20 ± 3 %, ns p > 0.05). We calculated the kinetics of fluorescence decay after stimulation using a one-phase exponential decay model (**Fig. 2F - H**). As no significant difference in the rate of the fluorescence decay was observed, we wondered if the pHuji/ChR2 combination works properly. One concern was that pHuji exhibits photo-switching behavior in blue light, as previously shown (Liu et al., 2021). This might explain the slight increase in fluorescence in animals without ATR after stimulation and could also influence the decay kinetics of animals with ATR (**Fig. 1F**). We thus investigated other possible protein combinations to achieve more accurate measurements of recycling kinetics using the pOpsicle assay.

**Fig. 2.**
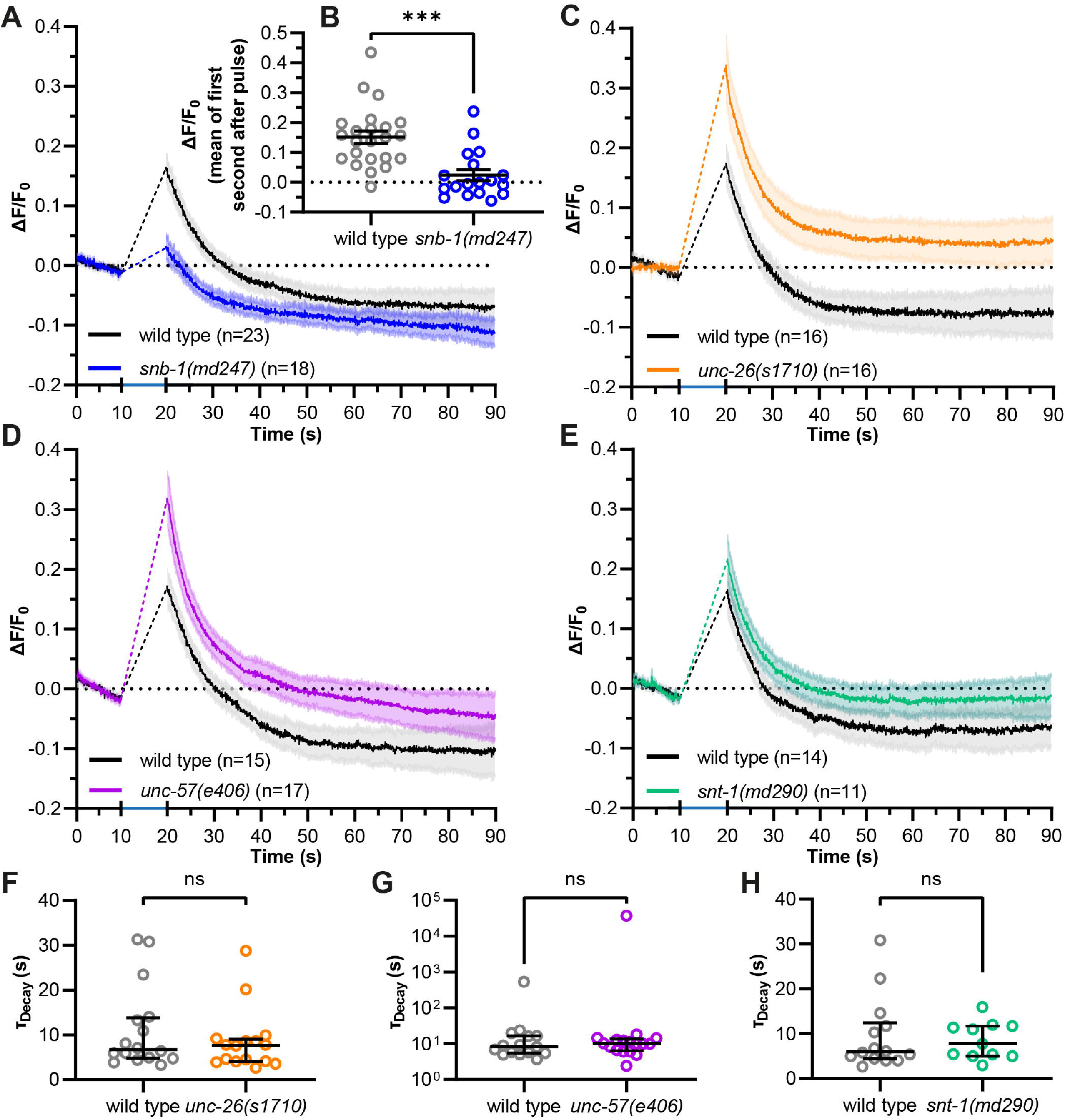
pHuji fluorescence signal increase is affected by mutation of *snb-1(md247)* in pOpsicle assays. **(A, C, D, E)** Mean (± SEM) DNC fluorescence of wild type and mutant animals treated with ATR, and expressing ChR2 and SNG-1::pHuji in cholinergic motor neurons. A 10 s continuous light pulse (460 nm, 0.34 mW/mm^2^, indicated by blue bar) was applied after 10 s. Number of animals is indicated (n), accumulated from N = 3 biological replicates. **(B)** Fluorescent signal of individual wild type and *snb-1(md247)* animals, as analyzed in (A), immediately following the end of the stimulation (20 – 21 s). Mean (± SEM). Unpaired *t*-Test; statistically significant difference is indicated as ***p < 0.001. **(F - H)** Calculated fluorescence decay constants of single animals using a one-phase exponential fit after stimulation (20 – 90 s). Median with interquartile range. Mann-Whitney test (ns, not significant, p > 0.05). In C – H, only animals showing a decay of fluorescence after stimulation were taken into consideration (wild type: 45 of 47 animals, *unc-26(s1710)*: 16 of 18, *unc-57(e406)*: 17 of 17, *snt-1(md290)*: 11 of 13).

### 3.3 Improving pOpsicle through combination of pHluorin with ChrimsonSA

We tested the more commonly used green fluorescent pHluorin as a pH-sensitive fluorophore which moreover features brighter fluorescence (Miesenbock et al., 1998; Li et al., 2022). Since pHluorin has overlapping excitation spectra with ChR2, we needed to exchange ChR2 with a red light activated channelrhodopsin. The recently described ChrimsonSA (for **s**uper red-shifted and **a**ccelerated), which is a mutated variant (S169A) of Chrimson, seemed to be a suitable candidate (Oda et al., 2018). Previously, we could show that ChrimsonSA can be used to depolarize *C. elegans* motor neurons upon stimulation with red light (Seidenthal et al., 2022; **Fig. 3A**). Two copies of pHluorin were inserted into the second intraluminal loop of SNG-1 as multiple pHluorin insertions have been shown to increase the signal-to-noise ratio (Zhu et al., 2009). We co-expressed this fusion construct with ChrimsonSA in cholinergic motor neurons and observed basal pHluorin fluorescence throughout the cholinergic nervous system (**Fig. 3B**). SNG-1::pHluorin also localizes to neuronal extensions of DA and DB neurons innervating dorsal muscle cells (**Fig. 3C**). Fluorescent puncta in the DNC indicate an accumulation of pHluorin at NMJs. To show that this fluorescence is indeed pH-dependent, we generated neurons as primary cell cultures from dissociated *C. elegans* embryos. pHluorin fluorescence could be observed in neurite extensions and around nuclei (**Fig. S1A, B**). Buffers containing ammonium chloride (NH_4_Cl), can be used to increase the pH within SVs (Sankaranarayanan et al., 2000; Dittman and Kaplan, 2006). When exposed to such a buffer, pHluorin fluorescence in neurites rapidly increased (**Fig. 3D** and **Fig. S1C**). The same culture was then treated with a low pH buffer to quench the surface fraction of pHluorin (**Fig. 3D** and **Fig. S1D**). The fluorescence immediately decreased to levels below the basal fluorescence. Washing of cells with a neutral-pH buffer then slowly increased the fluorescence again, showing the dependency of the green fluorescence on the surrounding pH. Animals expressing pHluorin and ChrimsonSA showed light dependent contraction of muscle cells in red light when treated with ATR (**Fig. 3E**). This indicates that ChrimsonSA is functioning properly to depolarize cholinergic motor neurons. Next, we performed pOpsicle assays with pHluorin animals (‘green pOpsicle’). DNC fluorescence gradually increased by 21 ± 2 % upon continuous 590 nm stimulation in ATR-treated animals (**Fig. 3F – I**, see **Fig. S1E** for a representation of all experiments, and **Supplementary Movie 1)**. The signal reached a plateau after ~ 10 s, indicating that most SNG-1::pHluorin was externalized at this point. Animals without ATR showed no increase upon stimulation (0 ± 1 %, ***p < 0.001; **Fig. 3H, I**). The increase in fluorescence was especially high in synaptic puncta along the DNC, indicating locations of highly active SV release sites (**Fig. 3F, G**). Following the end of the stimulation, like in red pOpsicle experiments, the signal gradually decreased towards levels before stimulation. Cultured neurons expressing pHluorin also showed a rise in fluorescence when illuminated (and when treated with ATR) even though the signal size was lower and only a fraction of the neurons showed a significant increase (**Fig. S1F**). Only 18 of 52 neurons showed a strong response, in contrast to intact animals, where this was the case for each individual recording (**Fig. 3I** and **Fig. S1E**; for a definition of “strong response”, see *Methods*). Furthermore, we observed spontaneous pHluorin increases in some animals, independent of treatment with ATR or illumination with red light (**Fig. S1G**).

**Fig. 3.**
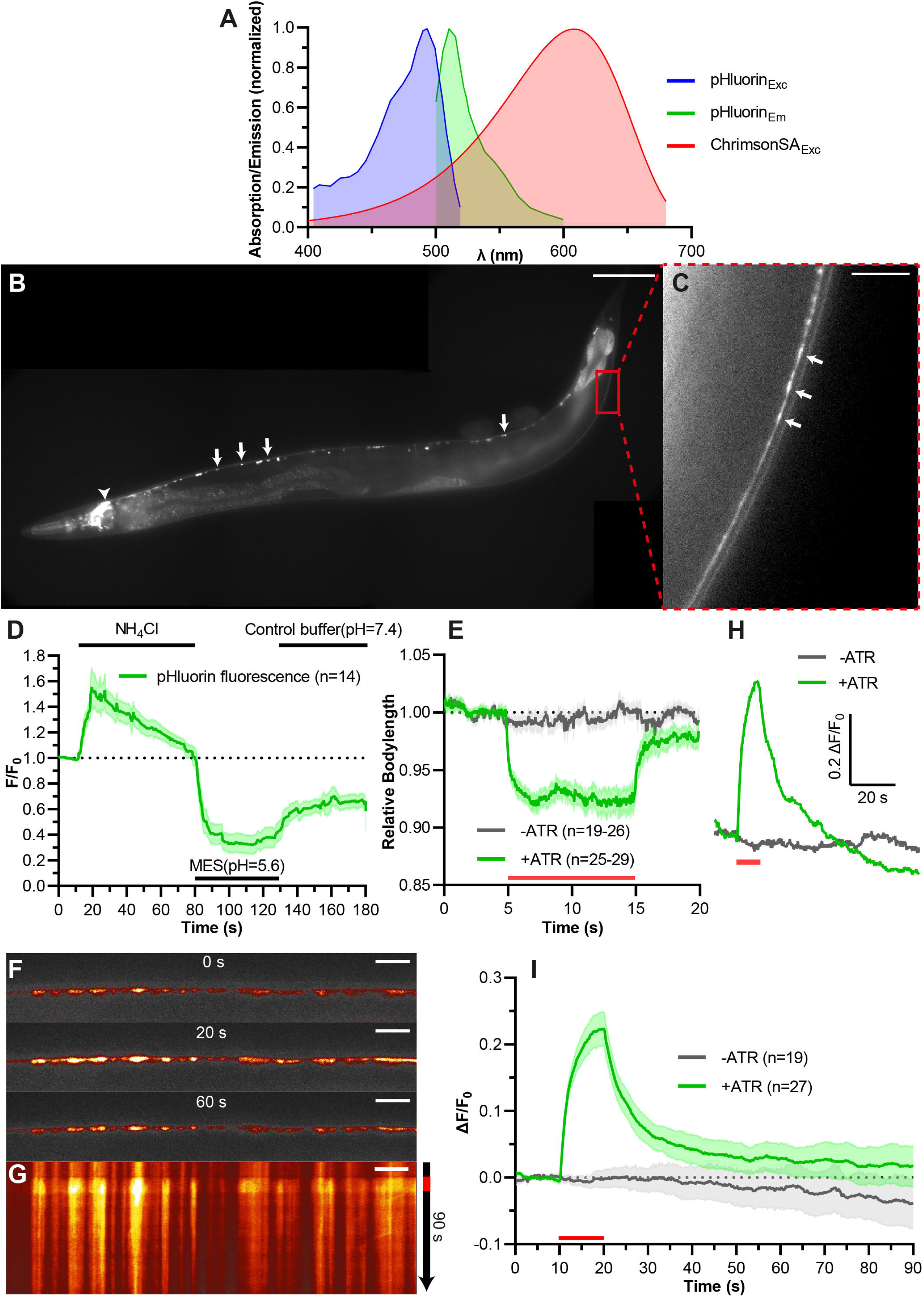
Combining ChrimsonSA with pHluorin (‘green pOpsicle’) for stimulation and visualization of exo- and endocytosis. **(A)** Relative excitation and emission spectra of ChrimsonSA and super-ecliptic pHluorin, normalized to the maximum absorption/emission amplitude. **(B)** Representative image of *C. elegans* expressing SNG-1::pHluorin in cholinergic neurons. Arrows: Cell bodies of A- and B-type motor neurons (ventral nerve cord). Arrowhead: cholinergic neurons in the head ganglia. 40 x magnification. Scale bar, 50 μm. **(C)** Enlarged view of the DNC in (B). Arrows: fluorescent puncta, representing en passant synaptic terminals. Scale bar, 10 μm. **(D)** Primary neuronal cell culture derived from *C. elegans* embryos, mean (± SEM) normalized change of pHluorin fluorescence in neurite extensions. NH_4_Cl containing solution (HEPES buffered, pH = 7.4) was added after 10 s. MES buffered solution (pH = 5.6) was added after 80 s. Cells were washed with control saline (HEPES buffered, pH = 7.4) after 125 s. Number of measured neurons is indicated (n), accumulated from N = 2 biological replicates. **(E)** Mean relative body length (± SEM) of animals expressing ChrimsonSA and pHluorin optionally treated with ATR, as indicated, normalized to the average body length before stimulation. A 10 s continuous light pulse (590 nm, 1 mW/mm^2^; indicated by red bar) was applied after 5 s. Number of animals (n), accumulated from N = 3 biological replicates. **(F)** Representative images of pHluorin fluorescence in the DNC of an animal treated with ATR at different time points during the pOpsicle assay, as indicated. A 10 s continuous light pulse (590 nm, 40 μW/mm^2^) was applied after 10 s. The ImageJ *Smart* Look-Up-Table was used. 100 x magnification. Scale bar, 5 μm. **(G)** Kymograph representing the change in fluorescence of the DNC represented in (F) over a time course of 90 s. The red bar indicates the period of light stimulus. Scale bar, 5 μm. **(H)** Representative traces of normalized DNC fluorescence of individual animals with or without ATR. A 10 s continuous light pulse (590 nm, 40 μW/mm^2^) was applied after 10 s (red bar). **(I)** Mean (± SEM) change in DNC fluorescence of animals supplemented with and without ATR. A 10 s continuous light pulse (590 nm, 40 μW/mm^2^) was applied after 10 s (red bar). Number of animals (n), accumulated from N = 4 (+ATR), and N = 3 (-ATR) biological replicates.

### 3.4 Quantification of SV exo- and endocytosis kinetics by the pHluorin pOpsicle assay

Next, we used green pOpsicle to analyze mutants affecting SV fusion (*snb-1(md247), snt-1(md290)*) and/or recycling (*unc-26(s1710), unc-57(e406), snt-1(md290)*) (**Fig. 4**). In *snb-1(md247)* mutants, the signal increase during stimulation was almost completely abolished (**Fig. 4A, B** and **Fig. S2A)**. Only 2 of 19 animals showed a relevant response (compared to 23/23 in wild type), demonstrating that the increase of fluorescence during stimulation reports on SV exocytosis. Synaptojanin-1 (*unc-26(s1710)*) mutant animals also exhibited a reduced signal compared to wild type (**Fig. 4C, D** and **Fig. S2B**). Thus, green pOpsicle faithfully revealed the expected depletion of SVs in *unc-26(s1710)* mutants (Harris et al., 2000), unlike pHuji that displayed a higher fluorescence after stimulation (**Fig. 2C**). To allow calculating the kinetics of fluorescence rise and decay, animals which showed no significant increase in fluorescence or no decay after stimulation had to be removed from analysis (**Fig. S3A**). We observed a significantly slower rise of fluorescence in *unc-26(s1710)* mutant compared to wild type animals, as could be seen by the increased time constants of the signal curves obtained from single animals, when fitted to a one-phase exponential association kinetic (τ_Rise, wild type_ = 2.2 s, τ_Rise, *unc-26*_ = 5.1 s; **Fig. 4E**). This could indicate an exocytosis defect, or it could be due to the reduced number of SVs that are available for fusion in the *unc-26(s1710)* mutant. The kinetics of the fluorescence decay after stimulation showed a strong reduction, as expected for the SV recycling mutant (**Fig. 4F, G**). Consequently, the calculated time constants of one-phase exponential decay were significantly increased in *unc-26(s1710)* mutants (τ_Decay, wild type_ = 16.0 s, τ_Decay, *unc-26*_ = 44.8 s), closely matching previous results measured in mammalian synaptojanin-1 knockout neurons (Watanabe et al., 2018). Wild type animals on the other hand showed decay time constants which are in the range of previous measurements in *C. elegans* sensory neurons, *Drosophila* motor neurons or mammalian hippocampal neurons (Ventimiglia and Bargmann, 2017; Yao et al., 2017; Li et al., 2022). Endophilin-1 (*unc-57(e406)*) mutants show similar trends as *unc-26(s1710)* with a significantly reduced signal and significantly slower association and decay kinetics compared to wild type (τ_Rise, wild type_ = 2.5 s, τ_Rise, *unc-57*_ = 3.1 s, τ_Decay, wild type_ = 10.7 s, τ_Decay, *unc-57*_ = 25.4 s; **Fig. 4H – L** and **Fig. S2C**). However, the *unc-57(e406)* mutant phenotype seemed to be less severe than in *unc-26(s1710)* mutants in all aspects, which supports previous findings using optogenetic stimulation combined with electron microscopy (Kittelmann et al., 2013). Synaptotagmin-1 (*snt-1(md290)*) mutants also displayed a smaller increase in fluorescence (**Fig. 4M, N** and **Fig. S2D**). The time constants of fluorescence rise are significantly increased in mutant animals (τ_Rise,wild type_ = 2.7 s, τ_Rise,snt-1_ = 5.7 s; **Fig. 4O**), demonstrating the role of SNT-1 in SV fusion (Liewald et al., 2008; Yu et al., 2013). We further observed a significantly delayed decrease of fluorescence after stimulation, indicating that SNT-1 is involved in SV recycling at *C. elegans* NMJs (Poskanzer et al., 2003; Mochida, 2022; τ_Decay wild type_ = 7.9 s, τ_Decay *snt-1*_ = 43.5s; **Fig. 4P, Q)**. Since the increase in fluorescent signal was generally lower in mutant animals, we wondered whether higher time constants of decay were caused by a lower activation of synaptic transmission rather than decreased SV recycling rates. We thus compared decay time constants of individual animals with the respective signal size. However, there was no significant correlation within any of the analyzed genotypes (**Fig. S3B – D**). Therefore, recycling rates likely do not depend on the amount of SV fusion in this assay.

**Fig. 4.**
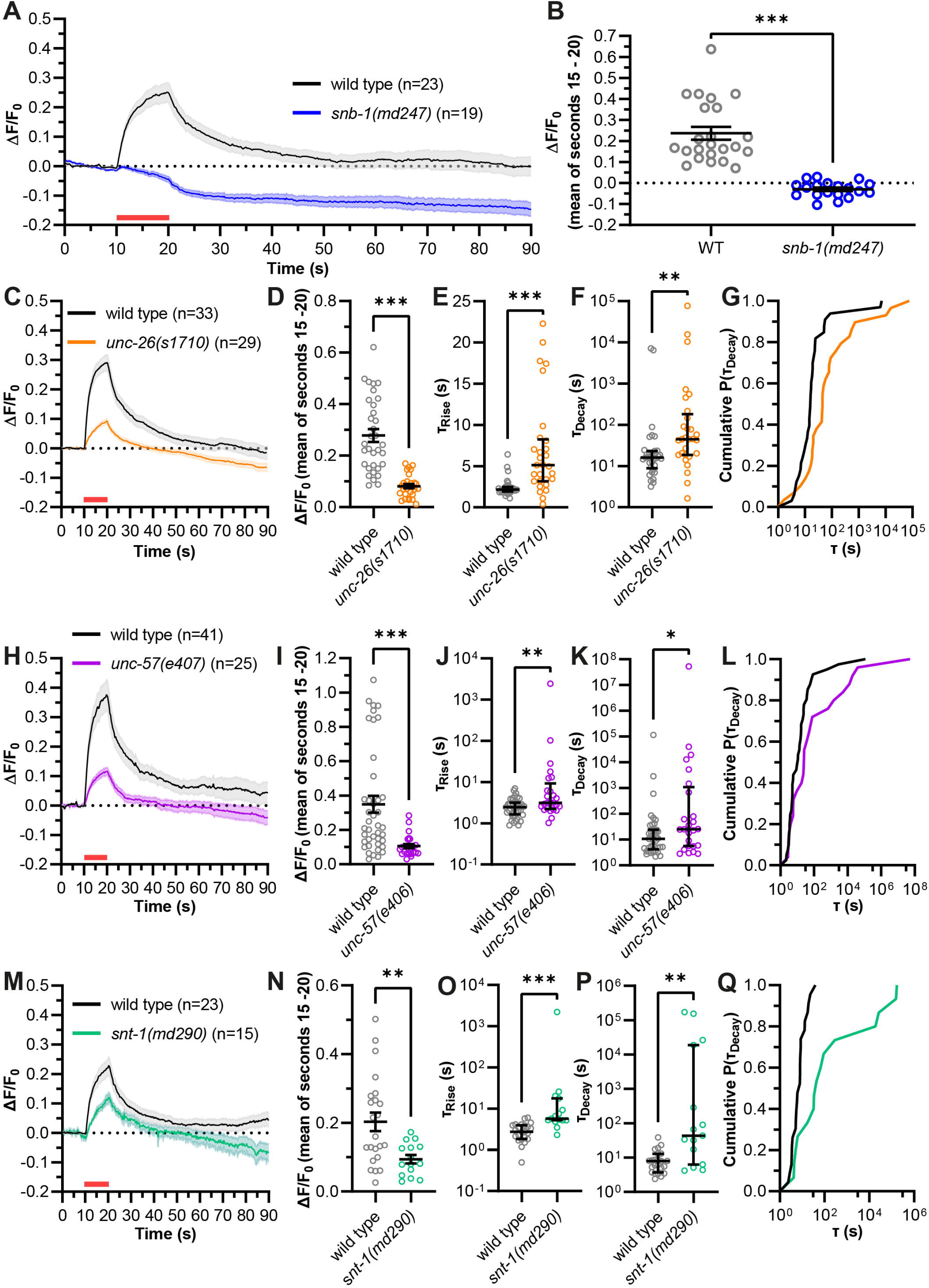
‘Green’ pOpsicle reports on mutations affecting SV fusion and endocytosis. **(A, C, H, M)** Mean (± SEM) change of fluorescence of SNG-1::pHluorin co-expressed with ChrimsonSA in cholinergic motor neurons. DNC of wild type and mutant animals, as indicated. A 10 s continuous light pulse (590 nm, 40 μW/mm^2^; indicated by a red bar) was applied after 10 s. Number of animals (n), accumulated from N = 4 - 6 biological replicates. **(B, D, I, N)** Fluorescent signal of individual wild type and mutant animals at the end of stimulation (15 – 20 s). Mean (± SEM). Unpaired *t*-Test (**p < 0.01, ***p < 0.001). **(E, J, O)** Calculated fluorescence rise constants of single animals using a one-phase exponential fit during stimulation (10 – 20 s). Median with interquartile range. Mann-Whitney test (**p < 0.01, ***p < 0.001). **(F, K, P)** Calculated fluorescence decay constants of single animals using a one-phase exponential fit after stimulation (20 – 90 s). Median with interquartile range. Mann-Whitney test (*p < 0.05, **p < 0.01). **(G, L, Q)** Cumulative frequency distribution of τ_Decay_ values displayed in (E), (I) or (M). In C – N, animals showing an increase < 3 standard deviations during stimulation, or no decay of fluorescence following stimulation, were excluded (wild type: 14 of 111 animals, *unc-26(s1710)*: 10 of 39, *unc-57(e406)*: 22 of 47, *snt-1(md290)*: 24 of 39).

### 3.5 Pulsed stimulation to potentially access different recycling mechanisms

Continuous photostimulation induces maximal depolarization and transmitter release, likely causing bulk endocytosis as the extreme form of ultrafast endocytosis (Kittelmann et al., 2013). To assess whether less vigorous, possibly more physiological activation also affects slower recycling, we applied 2 Hz pulsed stimulation (100 ms light pulses; **Fig. 5**). The mutant strains again showed significantly reduced signal amplitudes compared to wild type (**Fig. 5A, B** and **Fig. S4)**. *unc-26(s1710)* and *unc-57(e406)* mutants also showed significantly increased time constants of fluorescence rise (τ_Rise, wild type_ = 5.4 s, τ_Rise, *unc-26*_ = 21.3 s, τ_Rise, *unc-57*_ = 10.1 s, τ_Rise, *snt-1*_ = 7.2 s; **Fig. 5C**). When comparing pulsed and continuous stimulation we observed a tendency towards decreased fluorescence amplitudes and increased rise time constants, indicating that pulsed stimulation leads to a reduced activation of neurotransmission (**Fig. S5A, B**). For the recycling phase, pulsed stimulation again resulted in significantly larger decay time constants in *unc-57(e406)* and *unc-26(s1710)* mutants compared to wild type (τ_Decay,wild type_ = 11.3 s, τ_Decay, *unc-26*_ = 51144 s, τ_Decay, *unc-57*_ = 203.9 s; **Fig. 5D, E**), however, *snt-1(md290)* mutant animals showed no significant difference (τ_Decay, *snt-1*_ = 79.3 s). Possibly, SNT-1 is dispensable for recycling at lower levels of stimulation. Decay time constants for pulsed stimulation were not significantly different than for continuous stimulation, apart from *unc-57* mutants (**Fig. S5C**).

**Fig. 5.**
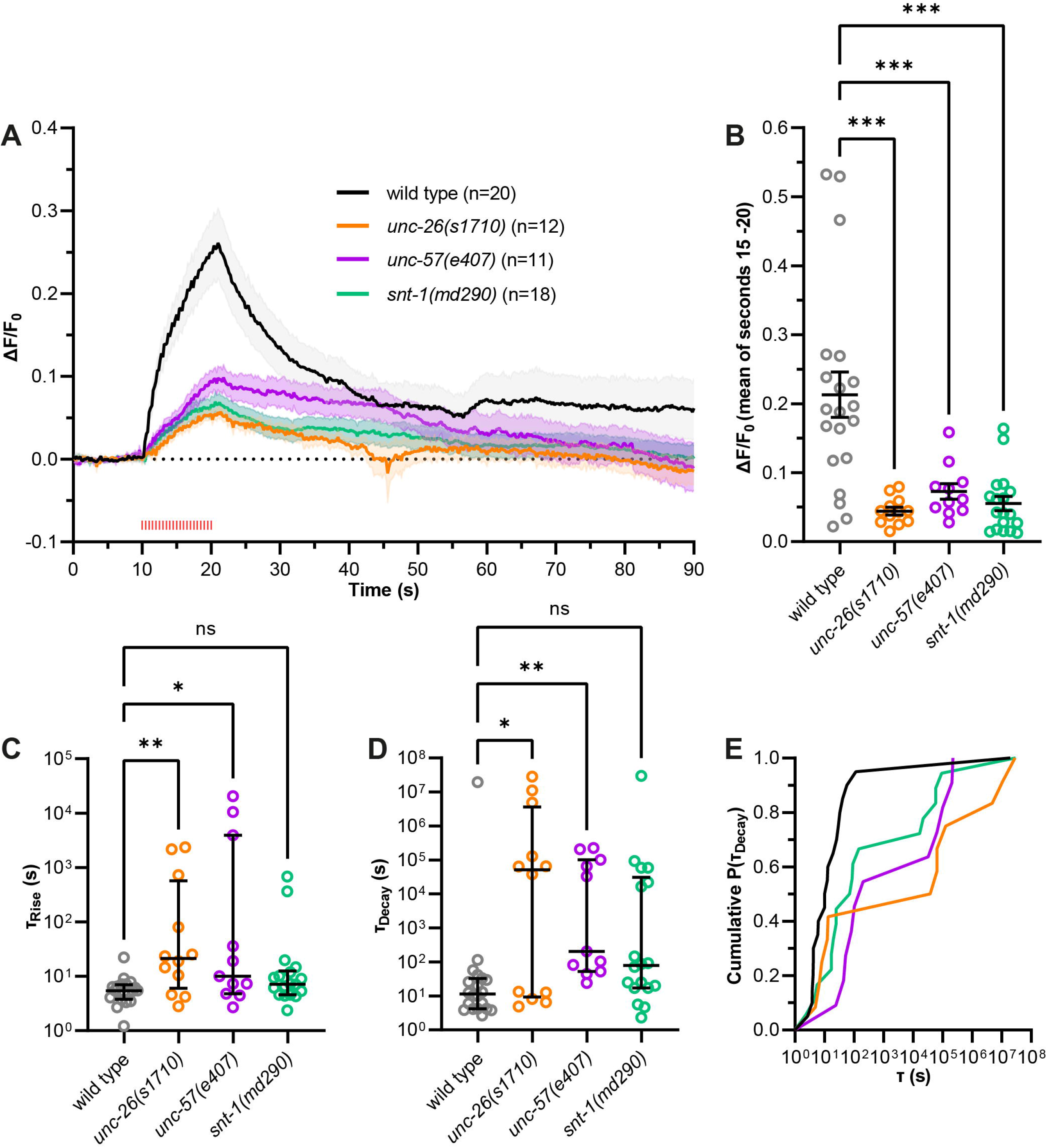
Using pulsed, more physiological optogenetic stimulation. **(A)** As in Fig. 4, but using 2 Hz pulsed light stimulation (100 ms pulses, 590 nm, 40 μW/mm^2^, red tick marks) was applied after 10 s for 10 s. Using this stimulation regime, more animals were excluded from analysis (wild type: 18 of 38 animals, *unc-26(s1710)*: 18 of 30, *unc-57(e406)*: 22 of 33, *snt-1(md290)*: 36 of 54) accumulated from N = 5 – 6 biological replicates. **(B)** Fluorescent signal of individual wild type and mutant animals at the end of stimulation (15 – 20 s). Mean (± SEM). One-way ANOVAs with Dunnett’s correction (***p < 0.001). **(C)** Calculated fluorescence rise constants of single animals using a one-phase exponential fit during stimulation (10 – 20 s). Median with interquartile range. Kruskal-Wallis test with Dunn’s correction (ns p > 0.05, *p < 0.05, **p < 0.01). **(D)** Calculated fluorescence decay constants of single animals using a one-phase exponential fit after stimulation (20 – 90 s). Median with interquartile range. Kruskal-Wallis test with Dunn’s correction (ns p > 0.05, *p < 0.05, **p < 0.01). **(E)** Cumulative frequency distribution of τ_Decay_ values displayed in (C).

### 3.6 pOpsicle reports on SV turnover in the single pair of RIM interneurons

While the green pOpsicle assay worked well in cholinergic motor neurons, it remained to be shown that this system works in other neuronal cell types. The RIM interneuron pair integrates signals from sensory neurons to regulate forward and reversal locomotion, using gap junctions as well as glutamate and tyramine signaling (Piggott et al., 2011; Li et al., 2020; Sordillo and Bargmann, 2021). We expressed SNG-1::pHluorin and ChrimsonSA in RIM using the *tdc-1* promotor. Green fluorescence could be observed in neuronal extensions surrounding the pharynx, suggesting correct localization of SNG-1::pHluorin (**Fig. 6A**). To explore ChrimsonSA functionality in this neuron pair, we measured animal crawling speed (**Fig. 6B**). Optogenetic depolarization of RIM neurons previously induced reversals and reduced crawling speed (Guo et al., 2009; Li et al., 2020; Sordillo and Bargmann, 2021). Consistently, illumination with red light slowed down crawling speed of RIM pOpsicle animals treated with ATR (**Fig. 6C**), while the number of reversals was increased. This indicated that ChrimsonSA depolarized RIM neurons when activated. Consequently, SNG-1::pHluorin fluorescence in synaptic puncta was significantly increased by ChrimsonSA activation (**Fig. 6D, E, Fig. S6** and **Supplementary Movie 2**), while control animals without ATR showed no change (+ATR 6 ± 2 %, -ATR −1 ± 1 %, ***p < 0.001; **Fig. 6F**). The signal increase was significantly slower than in cholinergic neurons (τ_Rise, RIM_ = 5.9 s, τ_Rise, cholinergic_ = 2.3 s; **Fig. 6G, H**), as was the decay of fluorescence following the end of stimulation (τ_Decay, RIM_ = 47.7 s, τ_Decay, cholinergic_ = 13.0 s; **Fig. 6I, J**). These results indicate that different classes of neurons may have diverging kinetics of SV exo- and endocytosis in *C. elegans*. Thus, the pOpsicle assay can be adapted to different neuronal cell types.

**Fig. 6.**
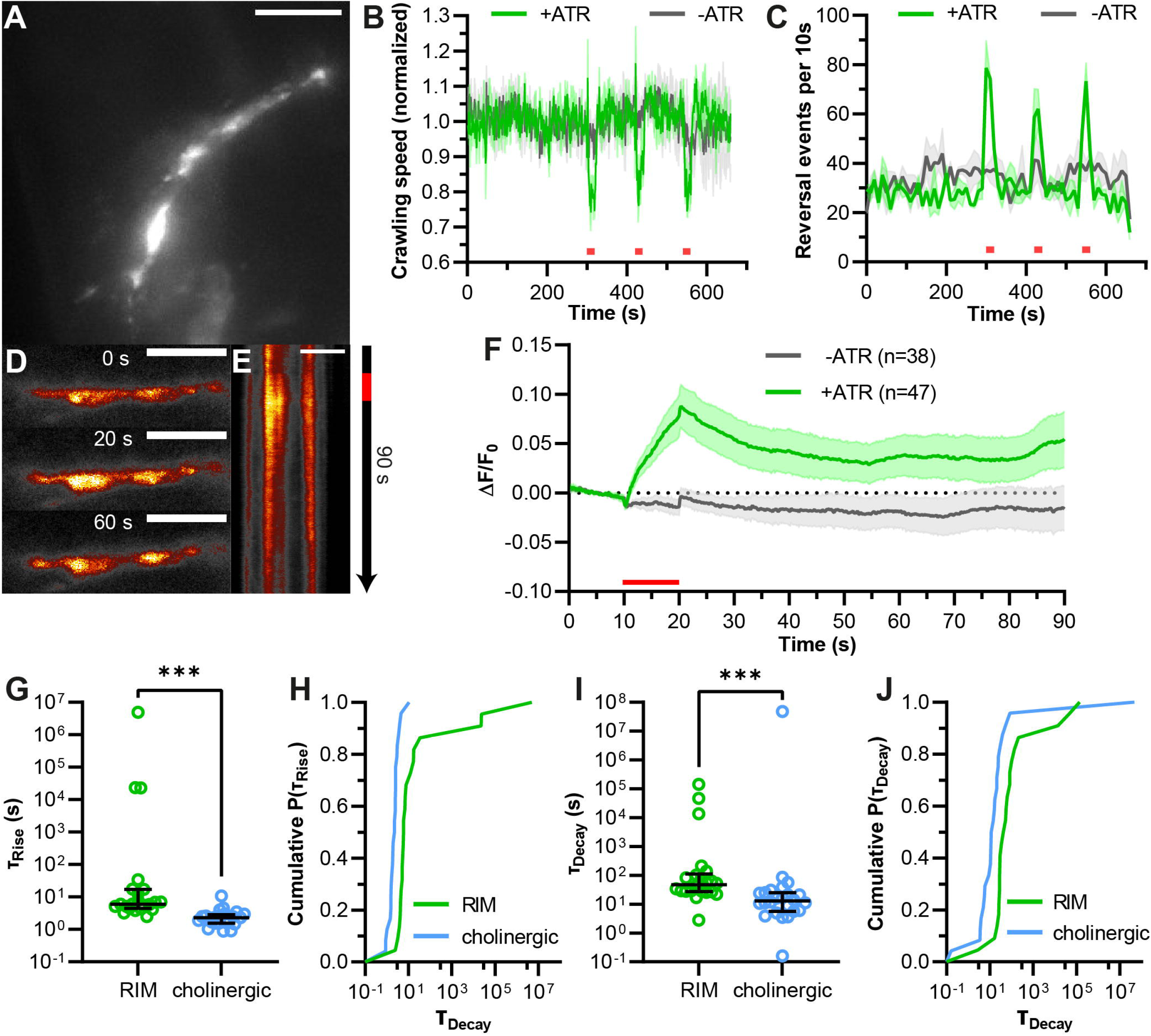
pOpsicle assay in the RIM interneuron pair. **(A)** Representative z-projected image of *C. elegans* expressing SNG-1::pHluorin in RIM neurons using the promotor of *tdc-1*. 40 x magnification. Scale bar, 5 μm. **(B)** Mean crawling speed (± SEM) of animals expressing ChrimsonSA and pHluorin in RIM neurons normalized to the average before the first light pulse. Three 20 s light pulses (623 nm, 400 μW/mm^2^) were applied at 300 s, 420 s, and 540 s as indicated by red bars. **(C)** Mean (± SEM) number of reversals in 10 s intervals. Light stimulation as in (B). In B, C, N = 3 populations of animals were tested. **(D)** Representative images acquired at different time points during the pOpsicle assay, pHluorin fluorescence in RIM neurons, animal treated with ATR. A 10 s continuous light pulse (590 nm, 40 μW/mm^2^) was applied after 10 s. The ImageJ *Smart* Look-Up-Table was used. 100 x magnification. Scale bar, 5 μm. **(E)** Kymograph representing the change in fluorescence in RIM neurons as shown in (D) over a time course of 90 s. The red bar indicates the period of light stimulus. Scale bar, 5 μm. **(F)** Mean (± SEM) pHluorin fluorescence in RIM neurons of animals with and without ATR. A 10 s continuous light pulse (590 nm, 40 μW/mm^2^) was applied after 10 s. Number of animals (n), accumulated from N = 5 (+ATR) or N = 4 (-ATR) biological replicates. **(G)** Comparison of fluorescence rise constants of single animals expressing pHluorin and ChrimsonSA in RIM neurons or in cholinergic neurons, using a one-phase exponential fit during stimulation (10 – 20 s). Median with interquartile range. Mann-Whitney test (***p < 0.001). **(H)** Cumulative frequency distribution of τ_Rise_ values displayed in (G). **(I)** Comparison of fluorescence decay constants of single animals as in G, using a one-phase exponential fit after stimulation (20 – 90 s). Median with interquartile range. Mann-Whitney test (***p < 0.001). **(J)** Cumulative frequency distribution of τ_Decay_ values displayed in (I). **(G-J)** Only animals showing a strong response during stimulation and a decay of fluorescence after stimulation were taken into consideration (RIM: 22 of 47 animals, cholinergic neurons: 24 of 27 animals as depicted in Fig. 3).

## 4 Discussion

Here, we present the first all-optical method to investigate SV recycling *in vivo* by combining pH-sensitive fluorescent proteins with ChRs. With pOpsicle, factors that influence the extent and rate of exo- and endocytosis can be investigated with minor experimental effort and equipment. We described two approaches using different pH-sensitive fluorescent proteins. pHuji could only be used to quantify the extent of exocytosis after stimulation. The low quantum yield and photoswitching behavior of this protein influenced the emitted fluorescence in a way that precluded quantification of exo- and endocytosis kinetics (Shen et al., 2014; Liu et al., 2021). Using pHluorin and ChrimsonSA, however, solved these problems, enabling calculation of fluorescence rise and decay time constants, which characterize different rates of exo- and endocytosis (Li et al., 2022). Neuronal primary culture of pHluorin expressing neurons could further open the way for investigation of exocytosis independent of SV recycling, e.g., by applying pharmacological agents such as bafilomycin A to inhibit SV acidification (Subramanian and Morozov, 2011; Li et al., 2022).

The pOpsicle system should be applicable to various neuronal cell types with minor modifications as exemplified by expression in cholinergic neurons and RIM interneurons. Establishing pOpsicle in various cell types could unveil disparities in SV exo- and endocytosis between different neuron classes, as has been shown for sensory neurons in *C. elegans* (Ventimiglia and Bargmann, 2017). Hitherto, electrophysiological recordings or Ca^2+^ imaging in body wall muscles were the method of choice to quantify neurotransmitter release in *C. elegans* in a time-resolved manner (Liewald et al., 2008; Wabnig et al., 2015). However, these only report postsynaptic effects which might be altered by unrelated phenomena such as neurotransmitter-receptor upregulation (Hammond-Weinberger et al., 2020). By using pOpsicle, a direct observation of the presynaptic SV cycle was achieved. This way, we could observe slowed SV fusion and endocytosis in RIM interneurons compared to cholinergic motor neurons. The faster release and recycling in cholinergic neurons may be in line with their function in mediating locomotion, and the likely high SV turnover needed, while the slower release rate could be important to the dual role of RIM in regulating reversal behavior (Li et al., 2020; Sordillo and Bargmann, 2021). While RIM promotes reversals through activation of AVA and AVE neurons via gap junctions, glutamate signaling inhibits reversal probability by reducing the amplitude of Ca^2+^ spikes within AVA and AVE (Li et al., 2020). A delayed release of glutamate from RIM may thus promote a fast reaction to noxious stimuli by initiation of reversals. Observation of pHluorin dynamics in freely moving worms may solve this issue. However, we note that it was imperative for the assay to work that the animals were kept as immobile as possible.

With green pOpsicle, we could reveal differences in recycling kinetics of synaptojanin-1 and endophilin-1 knockout mutants between continuous and pulsed stimulation. Discrepancies in the rate of recycling may be caused by different degrees of stimulation which trigger distinct routes of SV retrieval (Watanabe and Boucrot, 2017). Activity-dependent bulk endocytosis (ADBE) after strong optogenetic stimulation was shown to occur in *unc-57(e406)* and *unc-26(s1710)* knockout mutants and may be the main pathway of retrieval after continuous stimulation (Kittelmann et al., 2013; Nicholson-Fish et al., 2016; Yu et al., 2018). Moderate, pulsed stimulation however may trigger other endocytic mechanisms such as ultrafast and clathrin-mediated endocytosis, which are dependent on endophilin and synaptojanin (Milosevic et al., 2011; Watanabe et al., 2018). *snt-1(md290)* mutants displayed more severe recycling defects after strong stimulation, indicating that it is dispensable for recycling at lower activity. This is in stark contrast to previous results in mammalian hippocampal neurons in which synaptotagmin-1 promotes slow small-scale endocytosis, while inhibiting bulk retrieval during sustained neurotransmission (Chen et al., 2022). Although a role of SNT-1 in the slow clathrin-mediated endocytosis is likely also in *C. elegans* and simply not efficiently reported by pOpsicle, a role in inhibition of bulk endocytosis might not be conserved between nematodes and mammals. Previously, *snt-1(md290)* mutants have shown earlier fatigue of postsynaptic currents during strong optogenetic depolarization of cholinergic motor neurons indicating a compensatory SV recycling defect during strong sustained neurotransmission (Liewald et al., 2008; Wabnig et al., 2015).

Finally, SNG-1::pHluorin can further be used as a sensor for spontaneous neuronal activity and SV endocytosis in *C. elegans*, independent of optogenetic stimulation (**Fig. S1F**). This opens the way for multiplexing with other fluorescent reporters of neuronal activity such as genetically-encoded Ca^2+^ or voltage indicators (Dreosti and Lagnado, 2011; Jackson and Burrone, 2016; Azimi Hashemi et al., 2019). The combination of simultaneous imaging of SV dynamics and membrane potential changes could help to unravel the complicated interplay of interneurons in the control of locomotion by differentiating between electrical and chemical transmission.

## 5 Author Contributions

MS, BJ and XZ created plasmids and generated strains. MS performed pOpsicle, cell culture and contraction assays. MW tested ChrimsonSA in cholinergic neurons. NS and NE generated and tested ChrimsonSA/pHluorin strains. NR generated primary neuronal cell cultures. MS, BJ and AG designed and coordinated the study. MS and AG wrote the manuscript. AG supervised the work. All authors read and approved the final manuscript.

## 7 Acknowledgments

We are indebted to Franziska Baumbach and Hans-Werner Müller for their expert technical assistance, and members of the Gottschalk group for critical comments regarding the manuscript. We thank Dennis Vettkötter for providing MWT software. We also thank Johannes Vierock for providing ChrimsonSA absorption spectra. We thank Maximilian Bach for advice on RIM experiments, and the *Caenorhabditis* Genetics Center (CGC), which is funded by the NIH Office of Research Infrastructure Programs (P40 OD010440), for providing strains.

## Footnotes

1 https://github.com/MariusSeidenthal/pHluorin_Imaging_Analysis

