## Supplementary Figures S1 - S6 for "pOpsicle: An all-optical reporter system for synaptic vesicle recycling combining pH-sensitive fluorescent proteins with optogenetic manipulation of neuronal activity"

### 1 Supplementary Data

**Supplementary Movie 1.** Representative video of the DNC in an animal expressing SNG-1::pHluorin and ChrimsonSA in cholinergic neurons treated with ATR. A 10 s continuous light pulse (590 nm, 40 µW/mm2) was applied after 10 s as represented by red dot. The ImageJ Smart Look-Up-Table was used. 100 x magnification. Scale bar, 5 µm.

**Supplementary Movie 2.** Representative video of RIM neurons in an animal expressing SNG-1::pHluorin and ChrimsonSA in cholinergic neurons treated with ATR. A 10 s continuous light pulse (590 nm, 40 µW/mm2) was applied after 10 s as represented by red dot. The ImageJ Smart Look-Up-Table was used. 100 x magnification. Scale bar, 5 µm.

### 2 Supplementary Figures and Tables

#### 2.1 Supplementary Figures

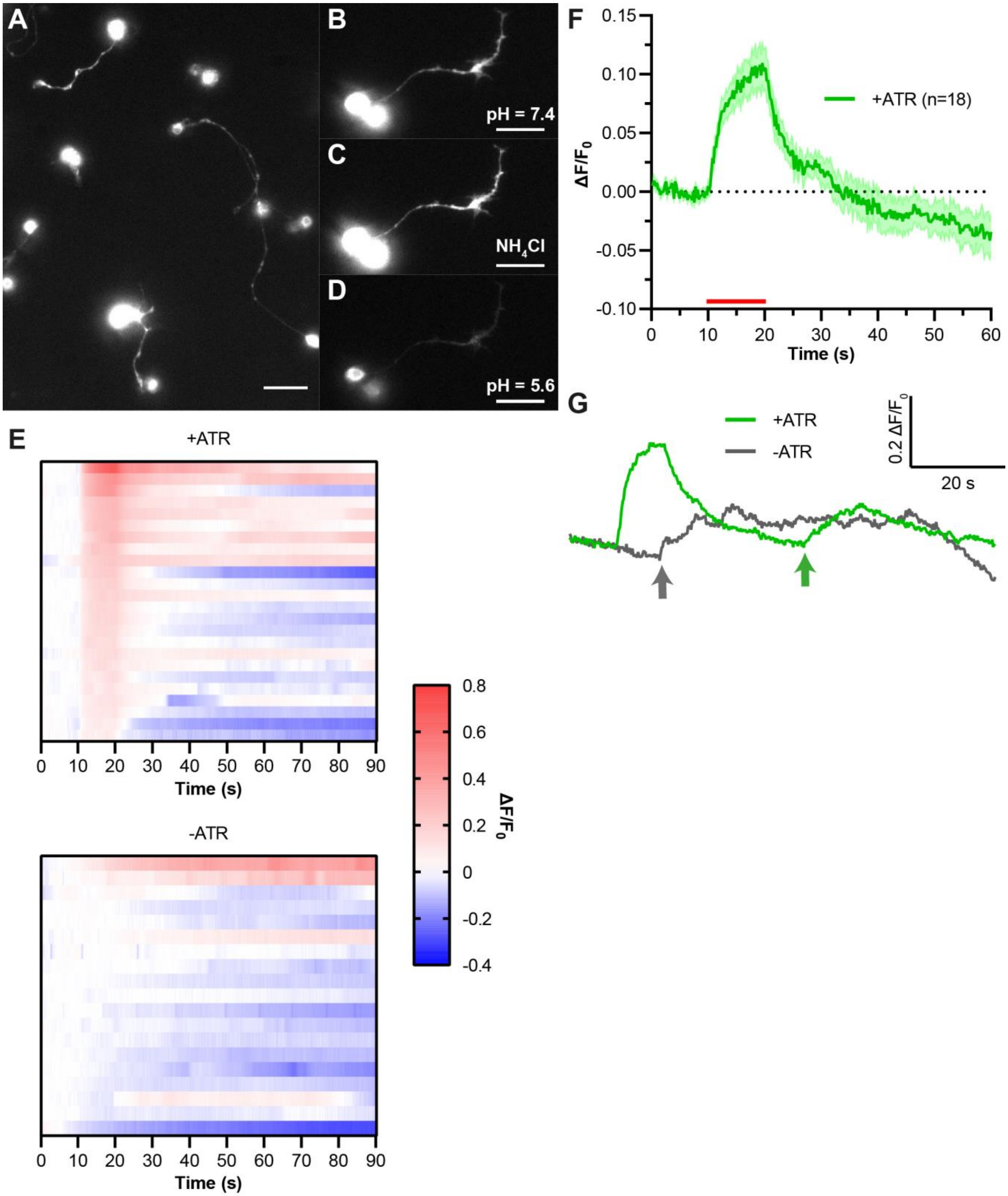
A
B
pH = 7.4
C
NH4Cl
D
pH = 5.6
E
+ATR
0 10 20 30 40 50 60 70 80 90
Time (s)
-ATR
0 10 20 30 40 50 60 70 80 90
Time (s)
0.8
0.6
0.4
0.2
0
-0.2
-0.4
ΔF/F0
F
0.15
0.10
0.05
0.00
-0.05
-0.10
ΔF/F0
+ATR (n=18)
0 10 20 30 40 50 60
Time (s)
G
+ATR
-ATR
0.2 ΔF/F0
20 s

**Figure S1. pHluorin imaging and pOpsicle assay in cultured primary cholinergic motor neurons.** **(A)** Representative image of primary neuronal cells expressing SNG-1::pHluorin. Scale bar, 10 µm. **(B, C, D)** Representative pHluorin expressing neuron, treated with different buffers as indicated above scale bars. Scale bar, 10 µm. **(E)** Color-coded traces representing normalized DNC fluorescence of individual animals represented in Figure 3H in the pOpsicle assay. **(F)** Mean ± SEM neurite fluorescence of pHluorin expressing cells in primary neuronal cell culture that have been supplemented with ATR. A 10 s continuous light pulse (590 nm, 40 µW/mm$^2$) was applied after 10 s. Only cells showing a strong response during stimulation were taken into consideration (18 of 52). **(G)** Exemplary pHluorin fluorescence traces recorded from animals with or without ATR, exhibiting spontaneous signal increases (indicated by arrows). A 10 s continuous light pulse (590 nm, 40 µW/mm$^2$) was applied after 10 s, triggering an increase of the pHluorin signal (indicated by arrows).

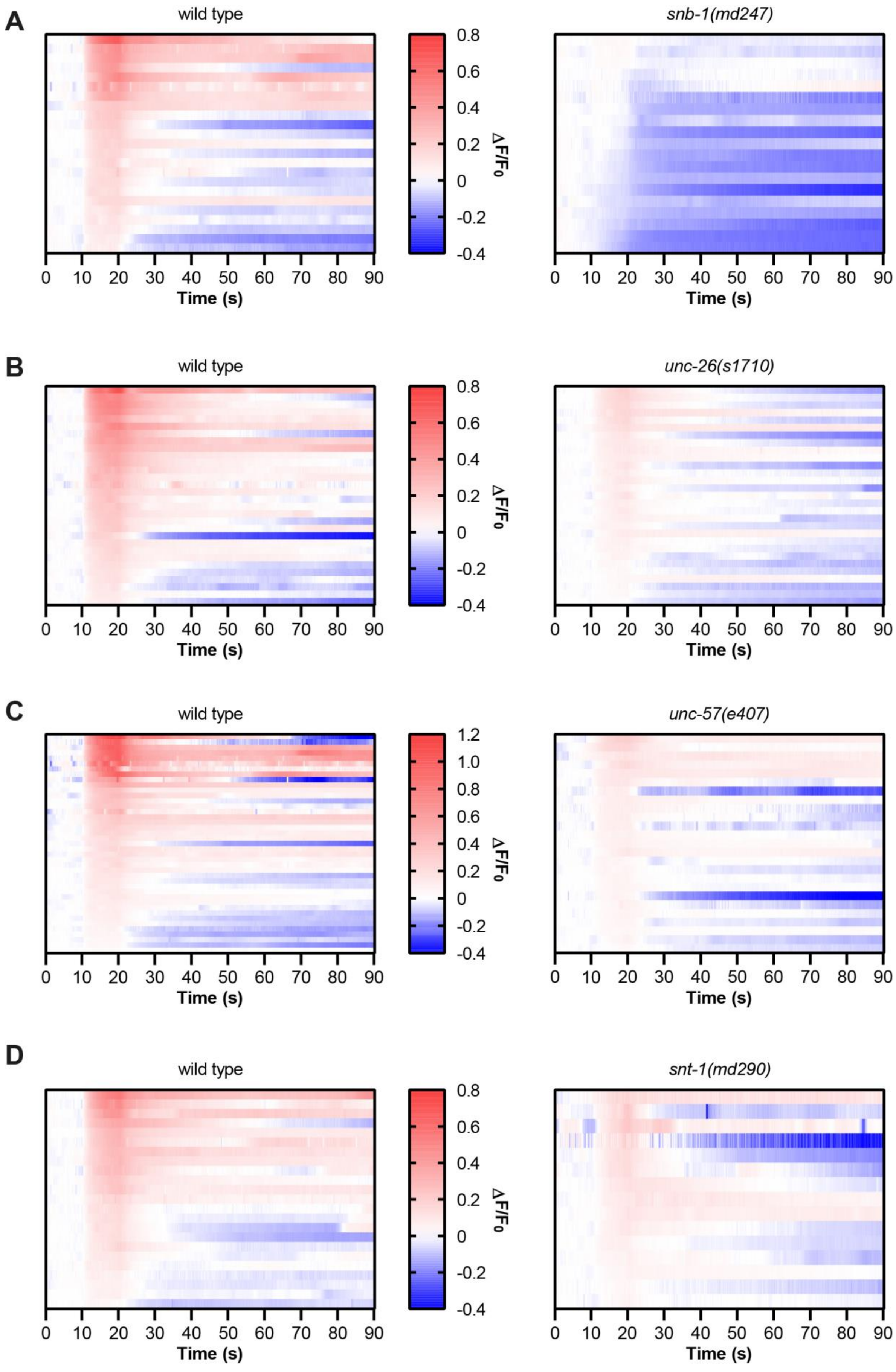
A
wild type
snb-1(md247)
B
wild type
unc-26(s1710)
C
wild type
unc-57(e407)
D
wild type
snt-1(md290)
ΔF/F0
0.8
0.6
0.4
0.2
0
-0.2
-0.4
1.2
1.0
0 10 20 30 40 50 60 70 80 90
Time (s)

**Figure S2. Individual traces of wildtype and mutant animals analyzed with the ‘green’ pOpsicle using continuous stimulation.** Color-coded traces representing normalized DNC fluorescence of individual animals depicted in Figures 4A, C, H and M in the pOpsicle assay.

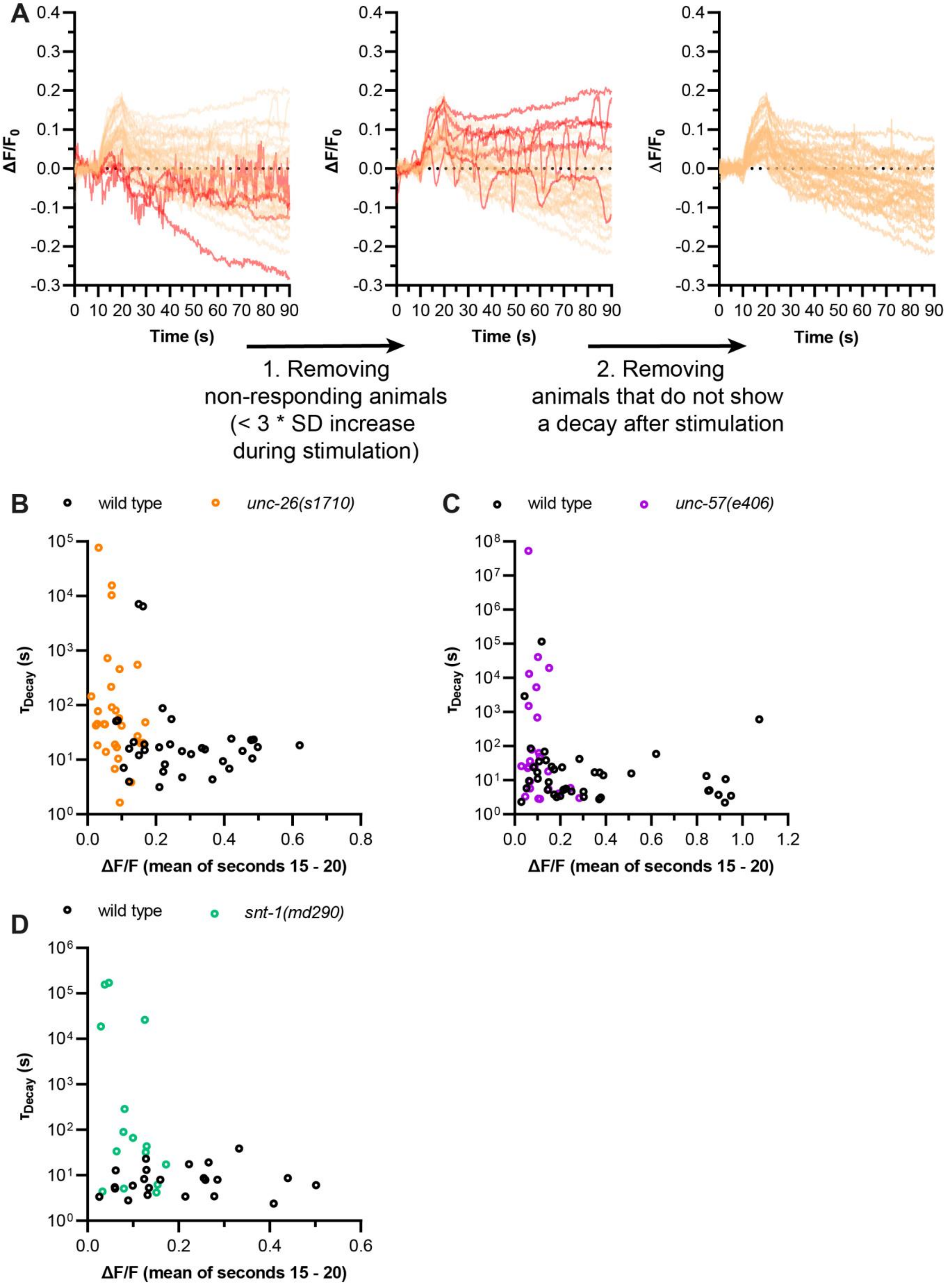
A
0.4
0.3
0.2
0.1
0.0
-0.1
-0.2
-0.3
ΔF/F0
0 10 20 30 40 50 60 70 80 90
Time (s)
1. Removing non-responding animals (< 3 * SD increase during stimulation)
2. Removing animals that do not show a decay after stimulation
B
wild type
unc-26(s1710)
τDecay (s)
ΔF/F (mean of seconds 15 - 20)
C
wild type
unc-57(e406)
D
wild type
snt-1(md290)

**Figure S3. Analysis of fluorescence decay time constants with the 'green' pOpsicle. (A)** Traces of individual *unc-26(s1710)* mutant animals as represented in Figure 4C are shown as an example of which datasets were removed for the analysis of fluorescence decay time constants. Removed dataset in each step are shown in red. Step 1: The maximum background corrected fluorescence during stimulation was calculated (as a moving average of 1 s). The animal was removed from analysis if this was lower than the average background corrected fluorescence before the stimulation + 3 * standard deviation (SD) of background corrected fluorescence before stimulation. Step 2: One-phase exponential decay fits after stimulation were inspected. If they showed an increase instead of decrease, the respective animal was discarded. **(B - D)** Fluorescence signals of individual wild type and mutant animals at the end of continuous stimulation (15 – 20 s) were compared to the respective calculated fluorescence decay constants. No significant correlation could be found for any of the datasets ($p > 0.05$). Spearman correlation was used.

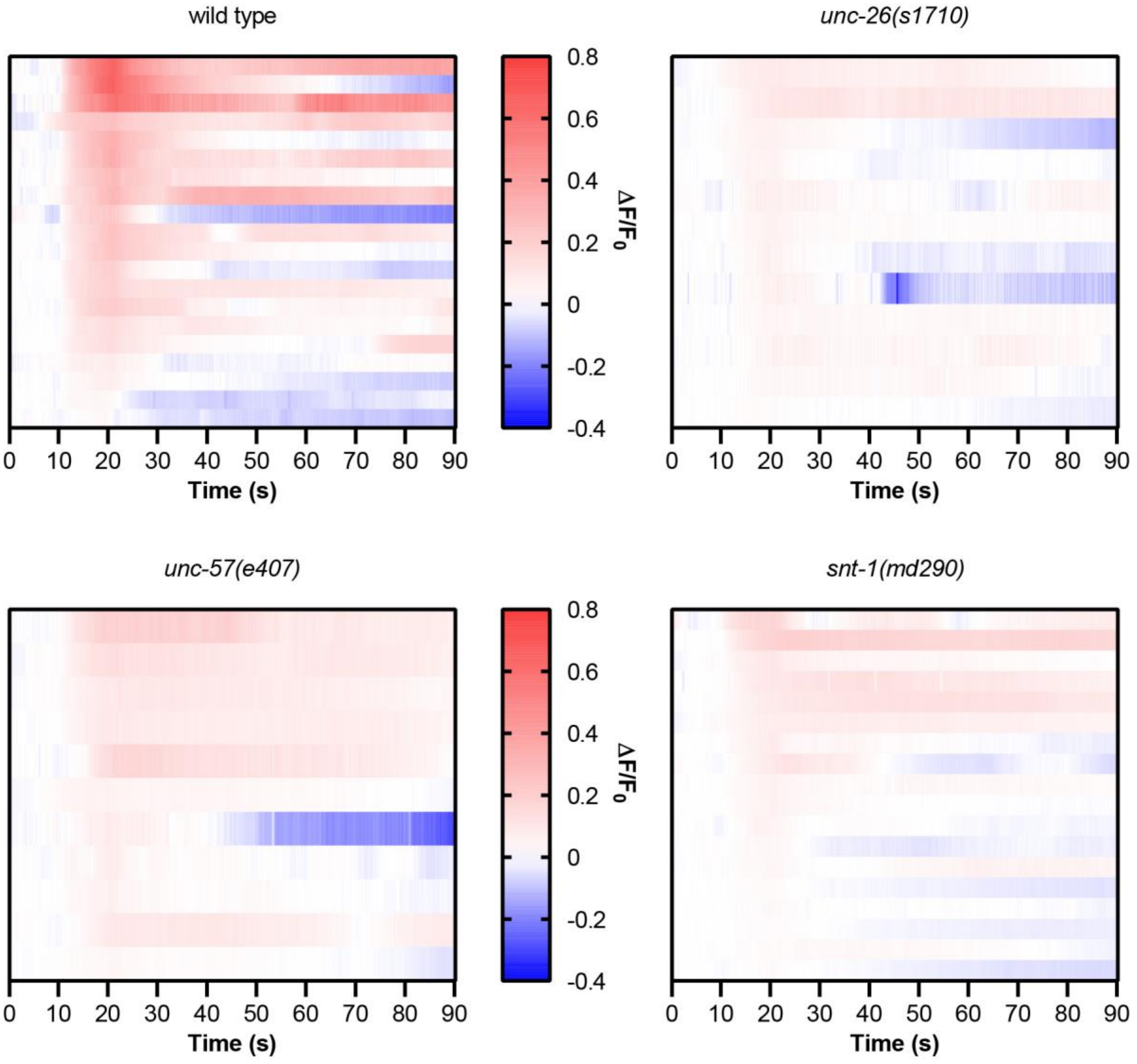

**Figure S4. Individual traces of wildtype and mutant animals analyzed with the 'green' pOpsicle using pulsed stimulation.** Color-coded traces representing normalized DNC fluorescence of individual animals depicted in Figure 5A in the pOpsicle assay.

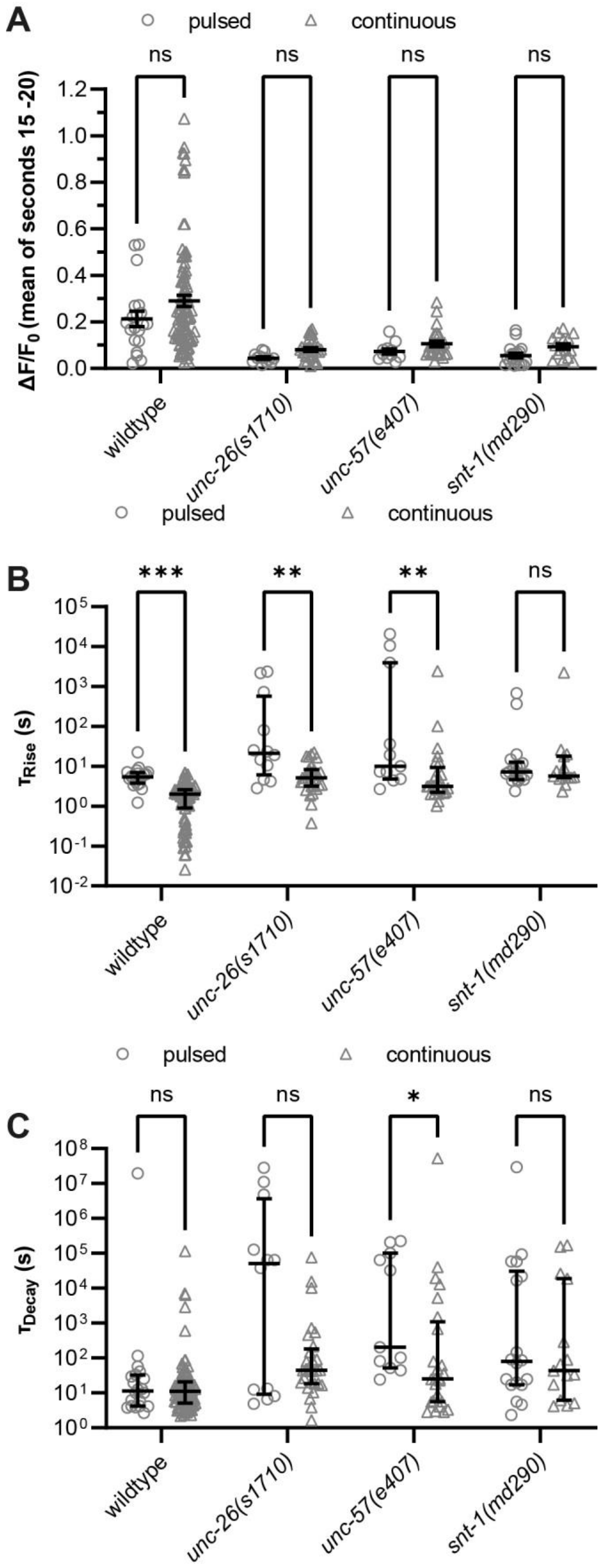
A
pulsed
continuous
ns
ns
ns
ns
ΔF/F0 (mean of seconds 15 -20)
1.2
1.0
0.8
0.6
0.4
0.2
0.0
wildtype
unc-26(s1710)
unc-57(e407)
snt-1(md290)
pulsed
continuous
B
***
**
**
ns
τRise (s)
10^5
10^4
10^3
10^2
10^1
10^0
10^-1
10^-2
wildtype
unc-26(s1710)
unc-57(e407)
snt-1(md290)
pulsed
continuous
C
ns
ns
*
ns
τDecay (s)
10^8
10^7
10^6
10^5
10^4
10^3
10^2
10^1
10^0
wildtype
unc-26(s1710)
unc-57(e407)
snt-1(md290)

**Figure S5. Comparison between continuous and pulsed stimulation. (A)** Fluorescent signal of individual wild type and mutant animals at the end of either pulsed or continuous stimulation (15 – 20 s). Mean (± SEM). Two-way ANOVA using Sidak's correction for multiple comparisons (ns, not significant). **(B)** Calculated fluorescence rise constants of single animals using a one-phase exponential fit during stimulation (10 – 20 s). Median with interquartile range. Multiple Mann-Whitney tests with Holm-Sidak correction for multiple comparisons (ns, not significant, $p > 0.05$, $^{**}p < 0.01$, $^{***}p < 0.001$). **(C)** Calculated fluorescence decay constants of single animals using a one-phase exponential fit after stimulation (20 – 90 s). Median with interquartile range. Mann-Whitney test (ns, not significant, $p > 0.05$, $^{*}p < 0.05$). Continuous: wild type n = 97 (pooled from animals depicted in Figure 4C – Q), *unc-26(s1710)* n = 29, *unc-57(e407)* n = 25, *snt-1(md290)* n = 15. Pulsed: wild type n = 20, *unc-26(s1710)* n = 12, *unc-57(e407)* n = 11, *snt-1(md290)* n = 18.

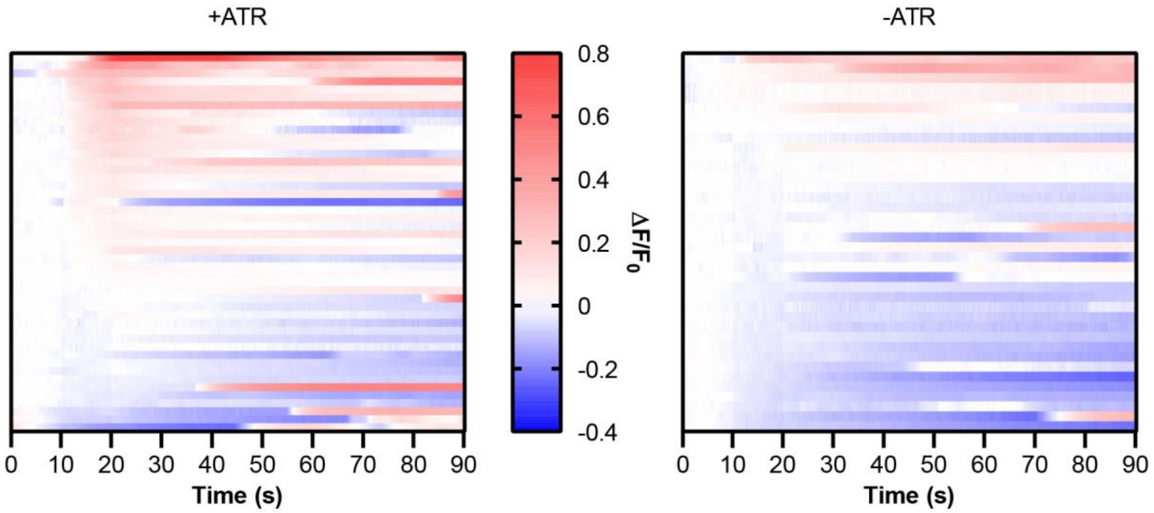


**Figure S6. Individual traces of the green pOpsicle assay in RIM neurons.** Color-coded traces representing normalized pHluorin fluorescence in RIM neurons of individual animals depicted in Figure 6F during the pOpsicle assay.
